# Co-option of neurotransmitter signaling for inter-organismal communication in *C. elegans*

**DOI:** 10.1101/275693

**Authors:** Christopher D. Chute, Elizabeth M. DiLoreto, Ying K. Zhang, Diego Rayes, Veronica L. Coyle, Hee June Choi, Mark J. Alkema, Frank C. Schroeder, Jagan Srinivasan

## Abstract

Biogenic amine neurotransmitters play a central role in metazoan biology, and both their chemical structures and cognate receptors are evolutionarily conserved. Their primary roles are in intra-organismal signaling, whereas biogenic amines are not normally recruited for communication between separate individuals. Here, we show that in *C. elegans*, a neurotransmitter-sensing G protein-coupled receptor, TYRA-2, is required for avoidance responses to osas#9, an ascaroside pheromone that incorporates the neurotransmitter octopamine. Neuronal ablation, cell-specific genetic rescue, and calcium imaging show that *tyra-2* expression in the nociceptive neuron ASH is necessary and sufficient to induce osas#9 avoidance. Ectopic expression in the AWA neuron, which is generally associated with attractive responses, reverses the response to osas#9, resulting in attraction instead of avoidance behavior, confirming that TYRA-2 partakes in sensing osas#9. The TYRA-2/osas#9 signaling system thus represents an inter-organismal communication channel that evolved via co-option of a neurotransmitter and its cognate receptor.

## Introduction

Inter-organismal communication occurs in several forms across the animal kingdom, both within and between species: prairie dogs use audio alarm calls to signal danger to conspecifics (1), birds display ornate visual cues and dances to attract mates (2), and honeybees dance to signal food location (3). Less apparent, though ancient and ubiquitous across all kingdoms of life, is chemical communication, which underlies social responses driven by chemosensation (4–7). Social chemical communication requires both intra- and inter-organismal signaling. First, a chemical cue is released into the environment by one organism that is then detected by specific receptors in another organism. Upon sensation, intra-organismal signaling pathways, e.g. neurotransmitter signaling, are activated that ultimately coordinate a social response.

Neurotransmitter monoamines such as dopamine, serotonin, tyramine and octopamine serve diverse functions across kingdoms (8). The associated signaling pathways often rely on highly regulated compound biosynthesis, translocation, either by way of diffusion or through active transport, and finally perception by dedicated chemoreceptors. Many neurotransmitters are perceived via G protein-coupled receptors (GPCRs); in fact, there appears to be a close relationship between GPCR diversification and neurotransmitter synthesis in shaping neuronal systems (9). Notably, the most common neurotransmitters share similar behavioral functions across phyla, for example, serotonin is commonly involved in regulating food responses (10–12). Other neurotransmitters, such as tyramine and octopamine, are only found in trace amounts in vertebrates, and in invertebrates act as adrenergic signaling compounds (13–15).

The nematode *Caenorhabditis elegans* affords many advantages for studying social chemical communication and neuronal signaling, namely, the animal’s tractability, well-characterized nervous system, and social behavioral responses to pheromones (16, 17). *C. elegans* secretes a class of small molecules, the ascaroside pheromones, which serve diverse functions in inter-organismal chemical signaling (18–20). As a core feature, these molecules include an ascarylose sugar attached to a fatty acid-derived side chain that can be optionally decorated with building blocks from other primary metabolic pathways (21). Ascaroside production, and thus the profile of relayed chemical messages, is strongly dependent on the animal’s sex, life stage, environment, and physiological state (22–25). Depending on their specific chemical structures and concentration, the effects of ascaroside signaling vary from social (e.g. attraction to icas#3) to developmental (e.g. induction of dauer by ascr#8) (Fig. 1A) (25–28). Furthermore, different combinations of these ascarosides can act synergistically to elicit a stronger behavioral response than one ascaroside alone, such as male attraction to ascr#2, ascr#3, and ascr#4 (19). Several GPCRs have been identified as chemoreceptors of ascaroside pheromones, such as SRX-43 in ASI in dwelling behavior and DAF-37 in ASK in hermaphrodite repulsion (29–33).

**Figure 1.**
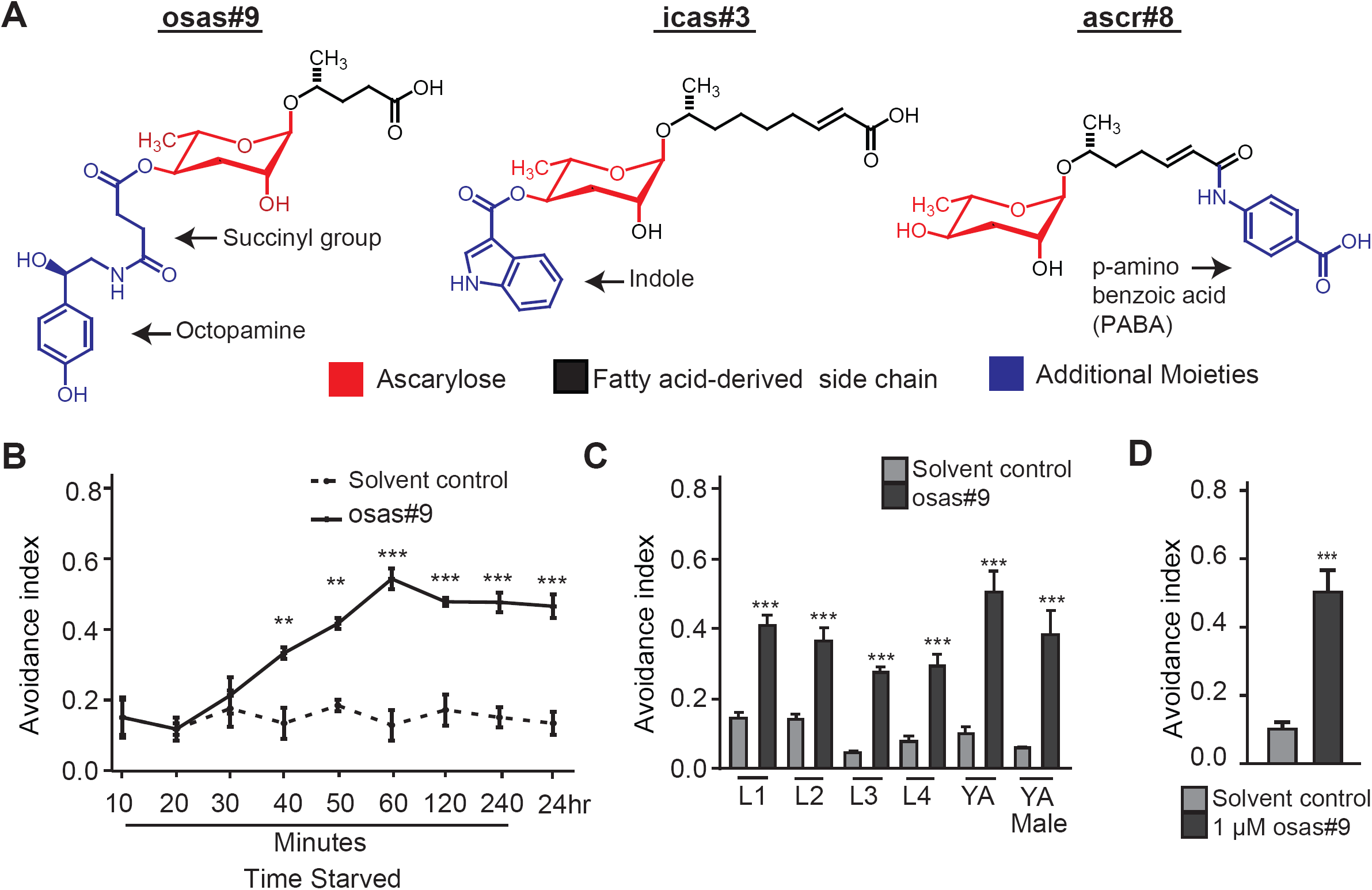
osas#9 is repulsive to starved animals. **A)** Structural and functional diversity of ascarosides. osas#9 is involved in avoidance, icas#3 attracts hermaphrodites and ascr#8 attracts males at low concentrations and induces dauer formation at high concentrations. **B)** Avoidance to osas#9 is dependent on the physiological state of *C. elegans*. Avoidance index of young adult (YA) wildtype (N2) animals in response to solvent control (SC) and 1 µM osas#9 after at different time points after removal from food. After 40 minutes of starvation, animals begin to avoid osas#9, and the response reaches a plateau at about 60 minutes, n ≥ 3 trials. Note for all other assays, unless otherwise stated, animals are starved for at least 60 minutes. **C)** All life stages of hermaphrodites and adult males avoid osas#9 when starved, n ≥ 4 trials. **D)** Avoidance index for starved young adult (YA) wildtype (N2) animals in response to the solvent control (SC) and to 1 µM osas#9, n = 8 trials. 1 µM osas#9 concentration was used in all other assays unless stated otherwise. Data presented as mean ± S.E.M; *P<0.05, **P<0.01, ***P<0.001, one factor ANOVA with Sidak’s multiple comparison posttest, except for Fig 1D, where student’s t-test was used. Asterisks displayed depict compared osas#9 avoidance response to respective solvent control.

Recently, an ascaroside, named osas#9, that incorporates the neurotransmitter octopamine was identified (22). Osas#9 is produced in large quantities specifically by starved L1 larvae and elicits aversive responses in starved, but not well fed conspecifics (22). The dependency on starvation of both its production and elicited response suggests osas#9 relays information on physiological status and unfavorable foraging conditions. However, it is unknown how osas#9 is perceived and drives starvation-dependent behavioral responses. Based on the unusual incorporation of a monoamine neurotransmitter building block in osas#9, we asked whether other components of monoamine signaling pathways have been recruited for inter-organismal signaling via osas#9. Here, we show that TYRA-2, an endogenous trace amine receptor, is required for the perception of osas#9, demonstrating co-option of a neurotransmitter and a neurotransmitter receptor for inter-organismal communication.

## Results

### Aversive responses to osas#9 require the GPCR TYRA-2

Previous work showed that production of the ascaroside osas#9 (Fig. 1A) is starkly increased in starved L1 larvae and elicits avoidance behavior in starved young adult hermaphrodites using a behavioral drop test assay (Fig. 1B) (22). This starvation dependent response is reversible: when animals are starved for an hour, and then reintroduced to food for two hours, no avoidance behavior is observed (Fig. S1A). For the current study we tested a broader range of conditions. We found that osas#9 elicits avoidance regardless of sex or developmental stage of animals (Fig. 1C), and that osas#9 is active over a broad range of concentrations (fM - µM) (Fig. S1B). 1 µM osas#9 was used for the remainder of this study unless otherwise noted (Fig. 1D). Ascarosides such as the male attractant ascr#3 and aggregation ascaroside icas#3 show activity profiles that are similarly broad as that of osas#9, whereas others, such as the mating cue ascr#8, are active only within more narrow concentration ranges (26, 34, 34).

The chemical structure of osas#9 is unusual in that it includes the neurotransmitter octopamine as a building block (Fig. 1A). Because octopamine and the biosynthetically related tyramine play important roles in orchestrating starvation responses, we investigated octopamine (*ser-3, ser-6, and octr-1*) and tyramine receptors (*tyra-2, tyra-3, ser-2, and ser-3*) for potential involvement in the osas#9 response (Fig. 2A) (36–40). We found that avoidance to osas#9 is largely abolished in a *tyra-2* loss-of-function (*lof*) mutant, whereas osas#9 avoidance was largely unaffected in the other tested neurotransmitter receptor mutants (Fig. 2A). We confirmed this phenotype was a result of the *lof* of *tyra-2* by testing a second *lof* allele of *tyra-2* (Fig. 2B), and by neuron-targeted RNAi (S2A,B) (41–43).

**Figure 2.**
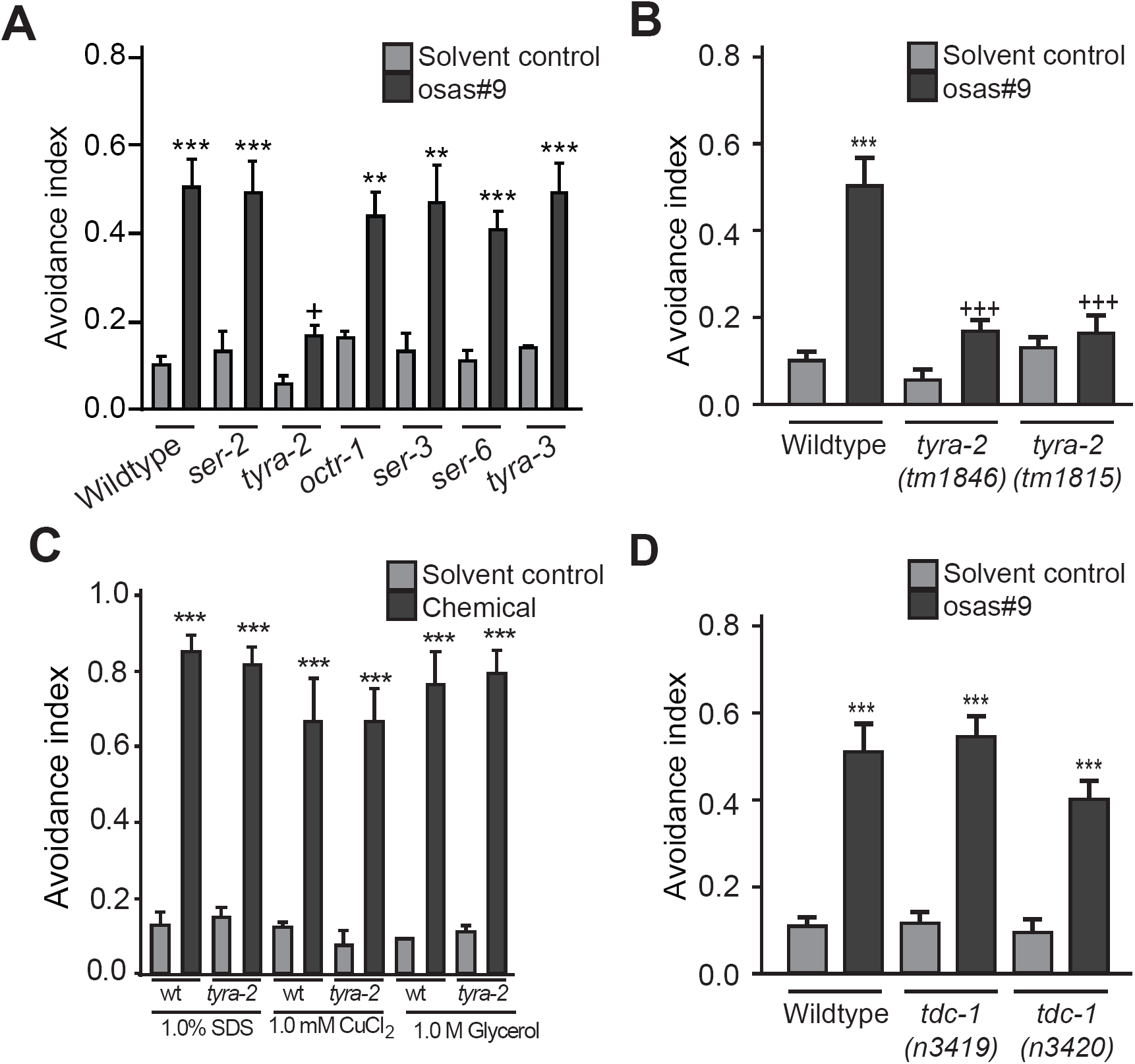
*tyra-2* is required for osas#9 aversive responses independent of tyramine. **A)** Screen for receptors required to mediate osas#9 avoidance. *tyra-2 lof* animals are defective in osas#9 avoidance response, n ≥ 4 trials. **B)** Two alleles of *tyra-2 lof* animals, *tm1846* and *tm1815*, are defective in osas#9 avoidance behavior, n ≥ 4 trials. *tyra-2*(*tm1846*) *lof* animals were used for the remainder of data presented in this manuscript. **C)** *tyra-2 lof* mutants showed no significant differences when subjected to known chemical deterrents, n ≥ 3 trials. **D)** osas#9 avoidance response is not dependent on endogenous tyramine. Two different alleles of *tdc-1 lof* animals, *n3419* and *n3420*, which lack tyramine biosynthesis, show normal response to osas#9, n ≥ 7 trials. Data presented as mean ± S.E.M; *P<0.05, **P<0.01, ***P<0.001, one factor ANOVA with Sidak’s multiple comparison posttest. Asterisks displayed without bar depict compared osas#9 avoidance to respective solvent control within groups. ‘+’ signs represent same p value as asterisks but representing difference between osas#9 avoidance of a strain/conditions in comparison to wildtype.

TYRA-2 is a G protein-coupled receptor (GPCR) that has been shown to bind tyramine with high affinity and octopamine to a lesser extent (38). To exclude the possibility that *tyra-2* is necessary for avoidance behaviors in general, we subjected *tyra-2 lof* animals to three well-studied chemical deterrents, SDS, copper chloride (CuCl_2_), and glycerol. No defects were found in the animals’ ability to respond aversively to these deterrents (Fig. 2C). This indicates that *tyra-2* is specifically required for osas#9 avoidance and is not part of a generalized unisensory avoidance response circuit. Since the response to osas#9 is dependent on physiological state, we examined whether *tyra-2* transcript levels changed under starved versus fed conditions using RT-qPCR. Starved animals exhibited a nearly two-fold increase in *tyra-2* expression (Fig. S2C).

We then asked whether tyramine signaling is required for the osas-9 avoidance response as *tyra-2* is known to bind to tyramine (38). We assayed two *tdc-1 lof* mutants, which lack the ability to synthesize tyramine (44). We observed that the behavioral response to osas#9 was unaltered in animals lacking tyramine biosynthesis (Fig. 2D). This demonstrates that the function of TYRA-2 in osas#9 avoidance is independent of tyramine, suggesting that TYRA-2 may be involved in perception of a ligand other than tyramine to promote aversive response to osas#9.

### tyra-2 is required in the ASH sensory neuron for physiological osas#9 response

We next asked where *tyra-2* is acting in the osas#9 aversion pathway. To determine the site of action of *tyra-2* in osas#9 avoidance, we designed a *tyra-2* translational fusion construct consisting of the entire genomic locus, including 2kb upstream, fused to GFP (p*tyra-2∷*TYRA-2*∷*GFP). We observed TYRA-2 expression in four sensory neurons: ASH, ASE, ASG, and ASI (Fig. 3A). These results are in agreement with previous expression studies on *tyra-2* localization (38) (Fig. 3A). We laser-ablated individual amphid sensory neurons to determine if a *tyra-2* expressing sensory neuron is required for the response. This revealed that ASH neurons are required for osas#9 response, whereas ablation of other neurons did not have a strong effect (Fig. 3B). We observed a slight reduction in the magnitude of the osas#9 aversive response in ASE-and ASI-laser-ablated animals (Fig. 3B); however, ASH/ASE and ASH/ASI double ablated animals did not differ in response from animals with ASH ablated alone, and ASE/ASI ablated animals did not differ from ASE or ASI alone (Fig. 3B). We then tested ASH, ASE, and ASI genetic ablation lines (45–48) and observed that at all tested concentrations, only ASH genetic ablation line resulted in complete abolishment of osas#9 avoidance (Fig. S3A,B,C). As with the laser ablation studies, we observed a slight decrease in osas#9 avoidance in ASE and ASI ablated animals (Fig. S3A,B,C) consistent with the findings for laser-ablated animals. Neurons not expressing *tyra-2* showed no defect in response to osas#9 (Fig. S3D). Our data implies that osas#9 is primarily sensed by ASH sensory neurons and that the ASE and ASI sensory neurons can potentially contribute by sensitizing ASH sensory neurons or by regulating downstream interneurons within the osas#9 response circuit.

**Figure 3.**
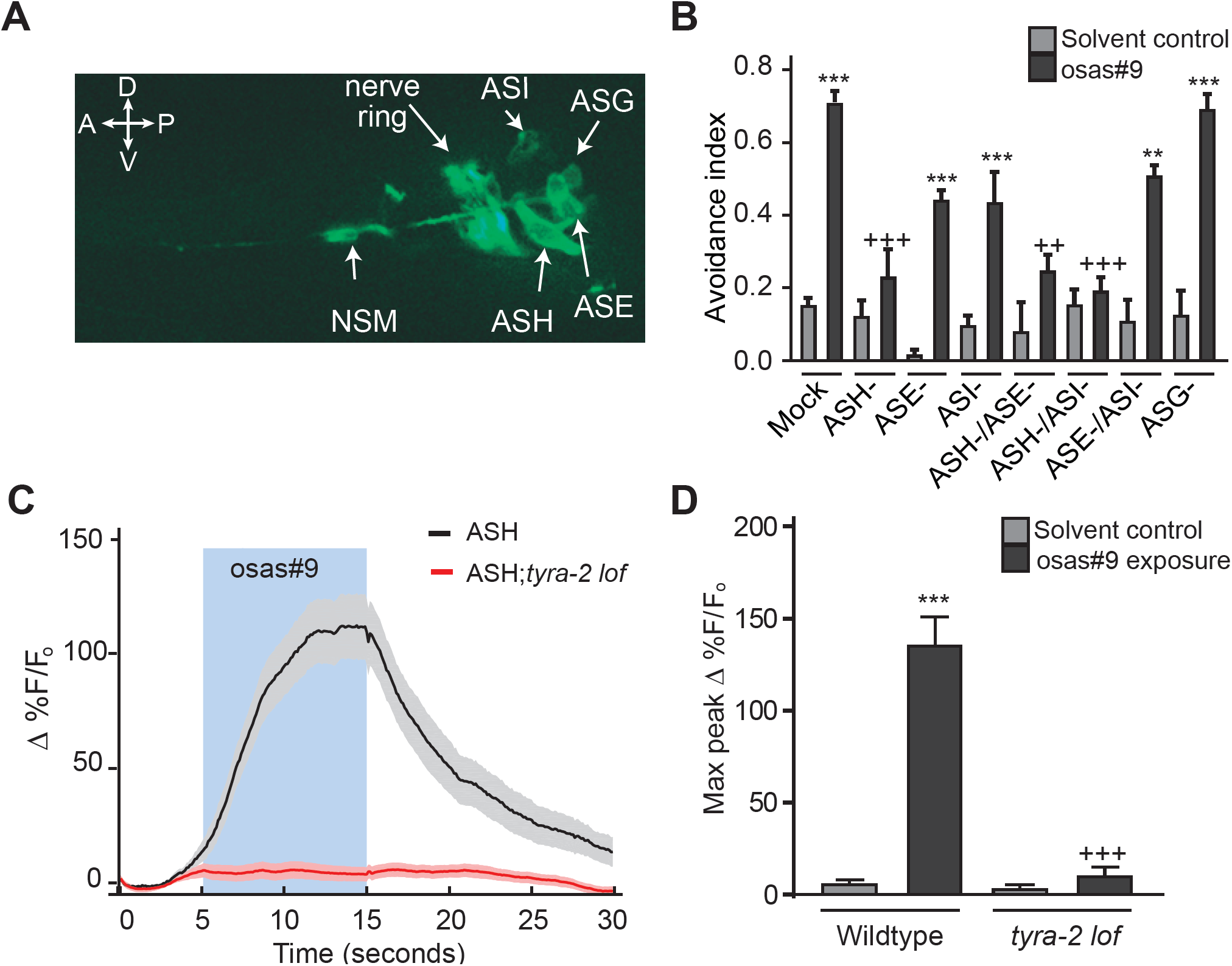
*tyra-2* expression in ASH sensory neurons is required for osas#9 response. **A)** Translational fusion consisting of 2kb upstream of the *tyra-2* gene and the entire *tyra-2* genomic locus was fused to GFP (*ptyra-*2∷*tyra-2∷*GFP) and injected in wildtype animals at 30 ng/µL revealing *tyra-2* expression in sensory neurons ASE, ASG, ASH, ASH, and NSM (40x magnification). **B)** Chemosensory neurons required for osas#9 response. Neurons expressing *tyra-2* reporter were ablated using laser microbeam. ASH neuronal ablations resulted in abolished response to osas#9 that was indistinguishable from solvent control. ASE and ASI ablated animals showed a reduced avoidance, but not to the extent of ASH neurons, n ≥ 3 trials with at least 10 ablated animals for each condition. **C,D)** Calcium dynamics of ASH neurons upon osas#9 exposure in a microfluidic olfactory chip. **C)** ASH∷GCaMP3 animals (black) display a change in calcium transients when exposed to osas#9. *tyra-2 lof* ASH∷GCaMP3 animals (red) did not display a change in fluorescence upon stimulation with the chemical. Shaded blue region depicts time when animals were subjected to the stimulus, n = 10 animals, 30 pulses. **D)** Maximum fluorescence intensity before (solvent control) and during exposure to 1 µM osas#9. Data presented as mean ± S.E.M; *P<0.05, **P<0.01, ***P<0.001, one factor ANOVA with Sidak’s multiple comparison posttest. Asterisks depict comparison between osas#9 and respective solvent control. ‘+’ signs represent same p value as asterisks but representing difference between osas#9 avoidance of a strain/conditions in comparison to wildtype.

To further elucidate the role of the ASH sensory neurons and TYRA-2 in osas#9 sensation, we utilized a microfluidic olfactory imaging chip that enables detection of calcium transients in sensory neurons (49, 50). We observed that, upon exposure to 1 µM osas#9, wildtype animals expressing GCaMP3 in the ASH sensory neurons exhibit robust increase in fluorescence upon stimulus exposure (Fig. 3C,D and Supplementary Video 1). Animals lacking *tyra-2* displayed no changes in fluorescence upon osas#9 exposure (Fig. 3C,D). These findings imply that *tyra-2* activity is necessary in ASH sensory neurons to sense and elicit osas#9 physiological responses.

Given that tyramine and octopamine are known ligands of TYRA-2, we also tested whether these neurotransmitters elicit aversive responses in *C. elegans* (38). Previous studies have shown that both tyramine and octopamine inhibit serotonin food-dependent increases in aversive responses to dilute octanol via specific G protein-coupled receptors (40). Both biogenic amines exhibited aversive behaviors at non-physiological concentrations much higher than required for osas#9, 1 mM for tyramine and octopamine compared to 1 µM for osas#9 (Fig. S4A,B, S1B). Similarly, high concentrations of tyramine (1mM) elicited calcium transients in ASH∷GCaMP3 but lower concentrations (1 µM) did not show calcium changes (Fig. S4C,D). Worms exposed to 1 mM octopamine displayed minimal change in calcium transients (Fig.S4C,D). These data show that the TYRA-2 receptor in the ASH sensory neurons is specifically involved in the avoidance response to osas#9. Tyramine or octopamine do not appear to be participating in the avoidance response, in agreement with the finding that tyramine biosynthesis is not required for avoidance to osas#9 (Fig. 2D).

### tyra-2 expression confers the ability to sense osas#9

Since expression of *tyra-2* in the ASH sensory neurons is required for calcium transients in response to osas#9, we asked whether *tyra-2* expression in the ASH neurons is sufficient to rescue the osas#9 behavioral response in *tyra-2 lof* animals. Expression of *tyra-2* under the *nhr-79* promoter, which is expressed in the ASH and ADL neurons, fully restored osas#9 avoidance (Fig. 4A,B) (51). To test whether expression of *tyra-2* in the ADL neurons is required for the phenotypic rescue, we ablated the ADL neurons in the transgenic animals. Ablation of the ADL neurons did not affect avoidance to osas#9 (Fig. 4C). Additionally, injection of the *tyra-2* translational reporter into *tyra-2* lof animals displayed sub-cellular localization in the ASH sensory cilia (Fig. 4D) and was observed to be functional as osas#9 aversion is rescued in these animals (Fig. 4E). These results affirm that the aversive behavioral response to osas#9 is dependent on *tyra-2* expression in the ASH neurons.

**Figure 4.**
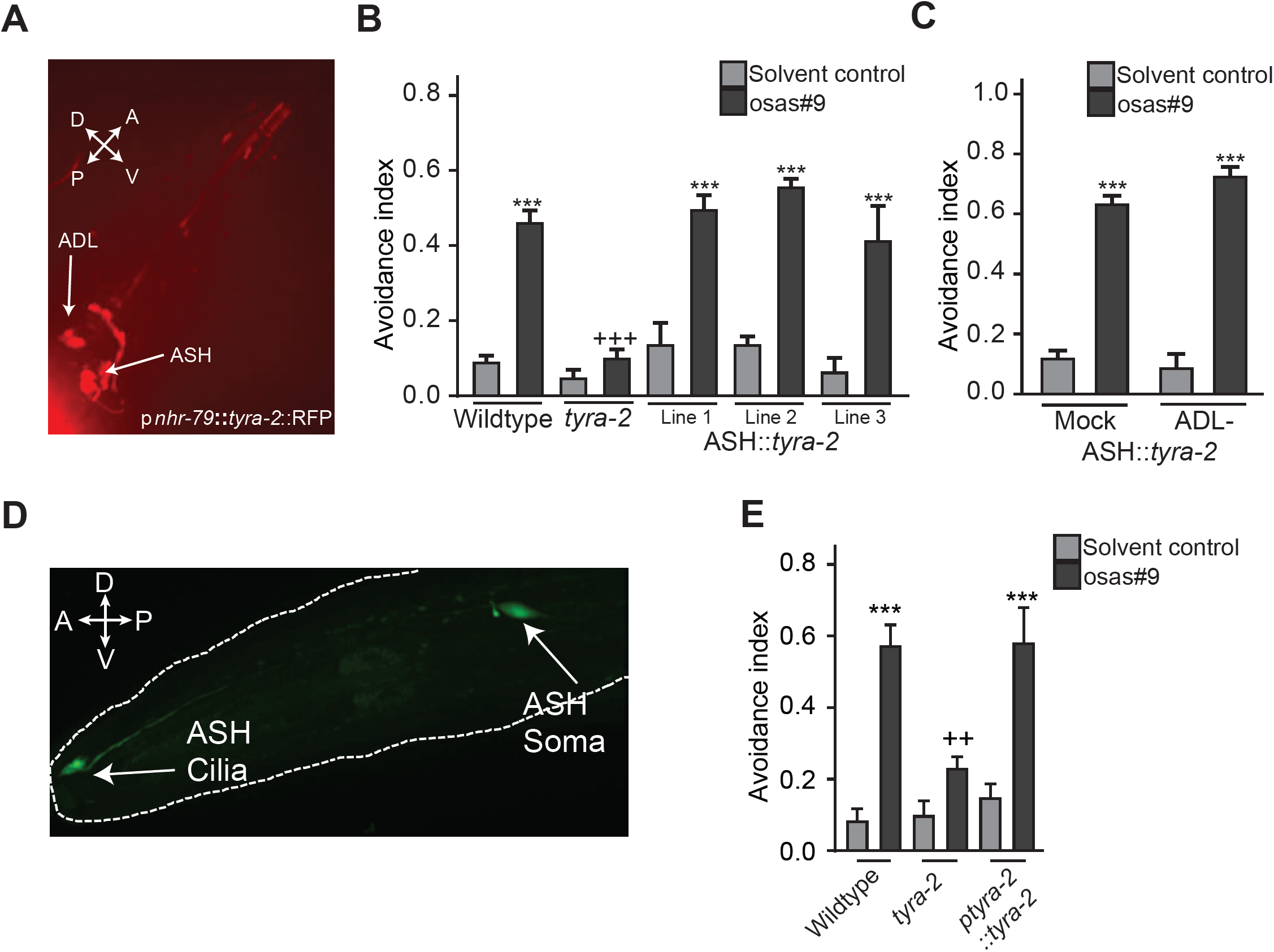
*tyra-2* expression is required in ASH sensory neurons for avoidance response to osas#9. **A)** A transcriptional rescue construct, p*nhr-79*∷*tyra-2*∷RFP exhibited expression of *tyra-2* in both ASH and ADL neurons (40x magnification). **B)** Rescue of *tyra-2* in ASH neurons fully reconstituted behavioral response to 1 µM osas#9, n ≥ 4 trials. **C)** Ablation of ADL neurons does not affect osas#9 avoidance in the rescue lines n≥4 trials. **D)** Sub cellular localization of *tyra-2*. A translational reporter of the entire *tyra-2* genomic locus (*ptyra-*2∷*tyra-2∷*GFP) was injected into *tyra-2 lof* animals at 1 ng/µL, revealing expression of the receptor in both soma and sensory cilia. (60x magnification). **E)** Expression of the translational reporter restores wildtype behavior in a *tyra-2 lof* background, n ≥ 5 trials. Data presented as mean ± S.E.M; *P<0.05, **P<0.01, ***P<0.001, one factor ANOVA with Sidak’s multiple comparison posttest. Asterisks depict comparison between osas#9 and respective solvent control. ‘+’ signs represent same p value as asterisks but representing difference between osas#9 avoidance of a strain/conditions in comparison to wildtype.

Previous studies in *C. elegans* indicate that behavioral responses (such as aversion or attraction) elicited by an odorant are specified by the olfactory neuron in which the receptor is activated in, rather than by the olfactory receptor itself (31, 52). Therefore, we asked whether expression of TYRA-2 in AWA neurons, which are generally involved in attractive responses to chemical cues (53, 54) would switch the behavioral valence of osas#9, resulting in attraction to osas#9, instead of aversion. Misexpression of *tyra-2* in the AWA sensory neurons in a *tyra-2 lof* background did not result in avoidance of osas#9, in contrast to expression of *tyra-2* in the ASH neurons (Fig. 5A). We then performed a leaving assay to test for attraction to osas#9 in the worms expressing *tyra-2* in the AWA neurons. This assay involves the placement of animals into the center of a NGM agar plate where osas#9 is present and measuring the distance of animals from the origin in one-minute intervals (Fig. 5B). *tyra-2 lof* animals displayed osas#9 leaving rates equal to the solvent control (Fig. 5C, S5), whereas worms misexpressing *tyra-2* in the AWA neurons displayed osas#9 leaving rates lower than that for solvent controls, indicating attraction (Fig. 5C, S5). Furthermore, worms misexpressing *tyra-2* in the AWA neurons stayed significantly closer to the origin than either wildtype or *tyra-2 lof* animals when exposed to osas#9 (Fig. 5C, S5). We confirmed that ectopic expression of *tyra-2* in AWA sensory neurons did not alter the native chemosensory parameters of AWA neurons (Fig. S6A,B). Hence misexpression of *tyra-2* in AWA neurons resulted in reprogramming of these nematodes, promoting attraction to the normally aversive compound osas#9.

**Figure 5.**
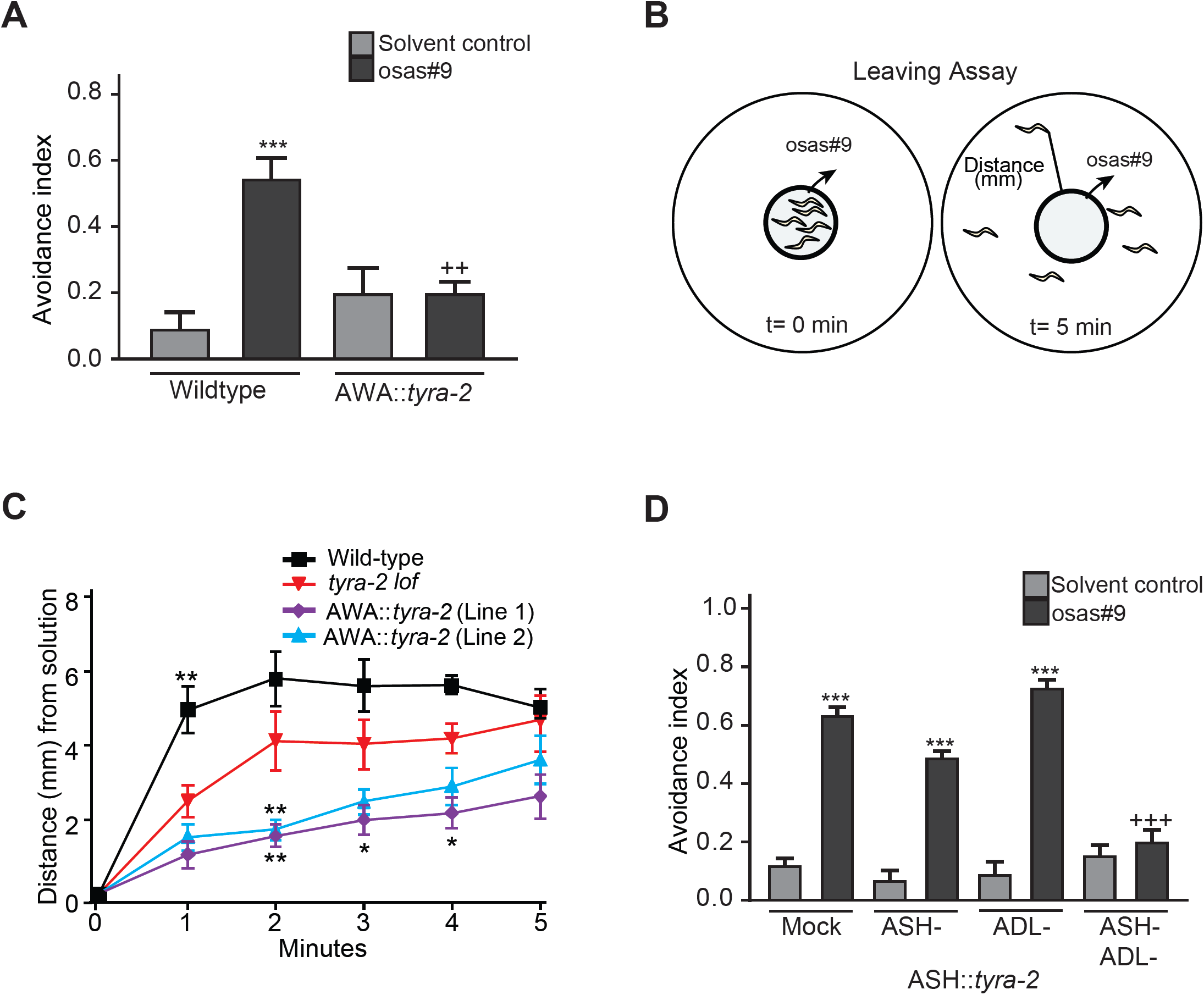
Ectopic expression of *tyra-2* confers the ability to respond to osas#9. **A)** Animals with reprogrammed AWA sensory neurons in *tyra-2 lof* background do not avoid osas#9, n ≥ 4 trials. **B)** Schematic illustration of the leaving assay to measure osas#9 attraction. (See material and methods for detailed description). **C)** Wildtype, *tyra-2 lof,* and AWA∷*tyra-2* lines were subjected to 10 pM osas#9 in the leaving assay. Wildtype animals left the osas#9 solution spot quicker than the *tyra-2 lof* animals, whereas the misexpression lines remained closer to osas#9, n ≥ 3 trials. **D)** Misexpression of *tyra-2* in ADL neurons confers avoidance behavior in response to osas#9. *nhr-79* promoter driving *tyra-2* in ASH and ADL sensory neurons rescues osas#9 avoidance. Ablation of ASH neurons in this line resulted in avoidance behavior to osas#9. Ablation of both ASH and ADL neurons in this line completely abolished avoidance, n ≥ 3 trials. Data presented as mean ± S.E.M; *P<0.05, **P<0.01, ***P<0.001, one factor ANOVA with Sidak’s multiple comparison posttest. Asterisks depict comparison between osas#9 and respective solvent control. ‘+’ signs represent same p value as asterisks but representing difference between osas#9 avoidance of a strain/conditions in comparison to wildtype.

Finally, we tested whether ectopic expression of *tyra-2* in the ADL neurons, which have been shown to detect aversive stimuli (55–58), results in a behavioral response to osas#9. For this purpose, we ablated the ASH neurons in the p*nhr-79*∷*tyra-2* strain, in which *tyra-2* is expressed in the ASH and ADL neurons. We found that these ASH ablated animals still avoid osas#9, similar to ADL ablated worms from this rescue line (Fig. 5D). Ablation of both the ASH and ADL neurons in this strain abolished the avoidance response (Fig. 5D). This implies that mis-expression of *tyra-2* in the ADL neurons confers the ability of this neuron to drive avoidance to osas#9. Taken together, results from both misexpression experiments (AWA and ADL neurons) demonstrate that TYRA-2 is necessary and sufficient to elicit osas#9-dependent behaviors.

### Gα protein gpa-6 is necessary in ASH sensory neurons for osas#9 avoidance

Since expression of the *tyra-2* GPCR is required in ASH neurons for osas#9 response, we sought to identify the Gα subunit necessary for osas#9 avoidance. Eight of the 21 Gα proteins are expressed in subsets of neurons that include the ASH sensory pair (*gpa-1, gpa-3, gpa-6, gpa-11, gpa-13, gpa-14, gpa-15,* and *odr-3*) (59–61). We tested mutants for each of those eight Gα subunits for their response to osas#9, (Fig. 6A) and found that *gpa-6 lof* animals do not avoid osas#9 (Fig. 6A). To determine whether *gpa-6* is necessary in ASH sensory neurons to mediate osas#9 responses, we expressed *gpa-6* using p*nhr-79* in the ASH neurons in a *gpa-6 lof* background. These animals displayed wildtype behavior when tested for osas#9 avoidance (Fig. 6B). To characterize cellular and sub-cellular localization of the *gpa-6* Gα subunit, we created a full length RFP translational fusion of the entire *gpa-6* locus including 4kb upstream. We detected *gpa-6* expression in the soma of AWA and ASH sensory neurons (Fig. 6C), in agreement with previous studies (60). However in addition to ASH soma localization, the translational fusion revealed presence of *gpa-6* in ASH cilia (Fig. 6C). Behavioral rescue by *gpa-6* expression specifically in the ASH neurons and its ciliary localization, support that this Gα subunit functions in mediating osas#9 avoidance.

**Figure 6.**
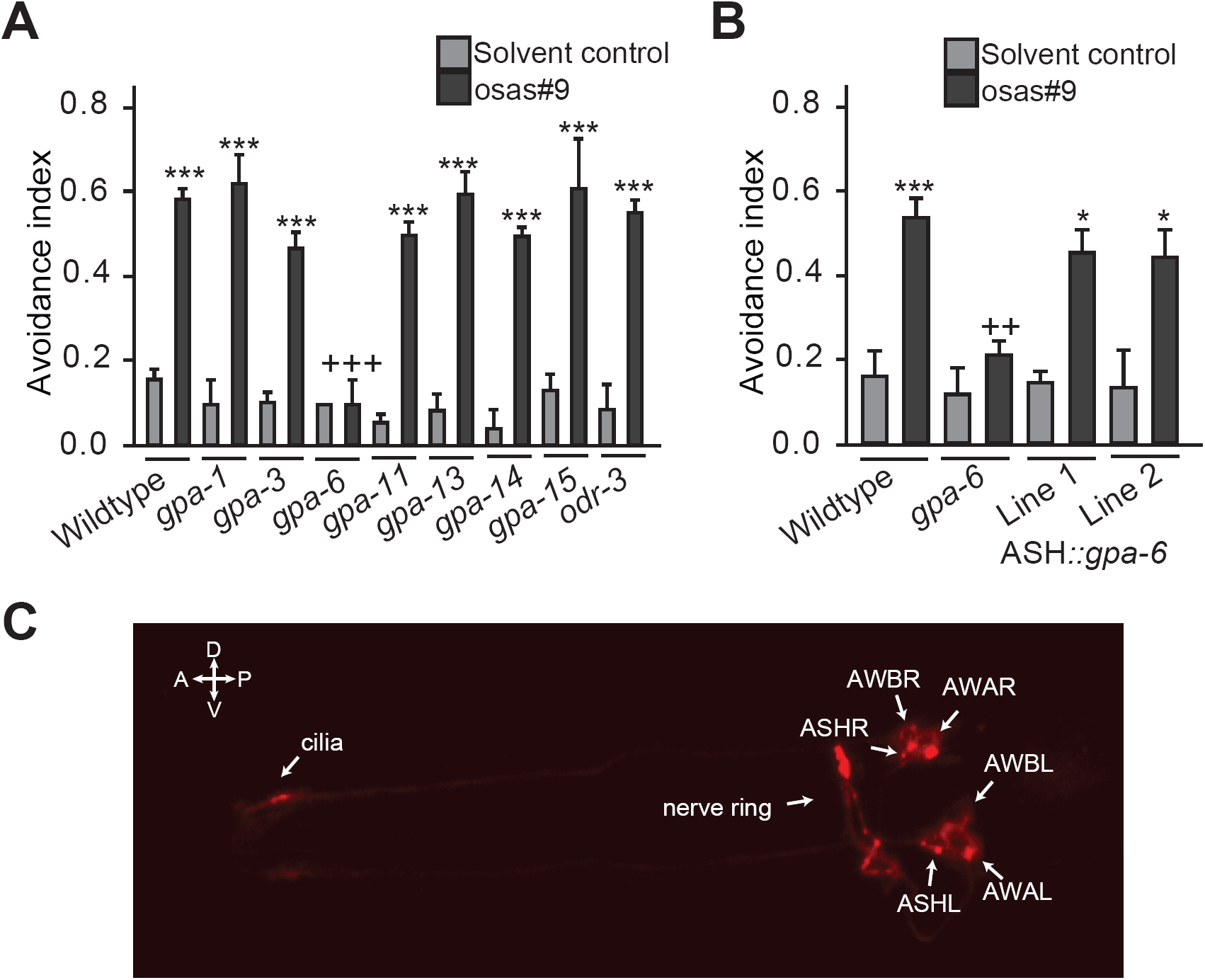
GPA-6 functions in ASH sensory neurons to mediate osas#9 response. **A)** Screen of mutations in G*α* subunits resulted in identification of the G*α*subunit *gpa-6*, which were defective in their avoidance response to osas#9, n ≥ 3 trials. **B)** Expression of *gpa-6* in ASH neurons using *nhr-79* promoter reconstituted avoidance response similar to wildtype animals, n ≥ 3 trials. **C)** *gpa-6* localizes to the soma and cilia in ASH neurons. Translational fusion of the entire *gpa-6* genomic region displayed localization of the subunit to the soma of AWA, AWB, and ASH neurons. In addition, we also observed ciliary localization in ASH neurons (40x magnification). Data presented as mean ± S.E.M; *P<0.05, **P<0.01, ***P<0.001, one factor ANOVA with Sidak’s multiple comparison posttest. Asterisks depict comparison between osas#9 and respective solvent control. ‘+’ signs represent same p value as asterisks but representing difference between osas#9 avoidance of a strain/conditions in comparison to wildtype.

## Discussion

How does a worm survive in changing environmental and physiological conditions? Given *C. elegans*’ complex ecology and a boom and bust lifestyle, worms need to make frequent adaptive developmental and physiological choices (62). The octopamine-derived pheromone osas#9, secreted in large quantities by L1 larvae under starvation conditions, appears to promote dispersal away from unfavorable conditions (Fig. 7). Here we show that this pheromone is detected by the GPCR *tyra-2*, a canonical neurotransmitter receptor that is expressed in the ASH sensory neurons. To our knowledge this is the first instance in which a “repurposed internal receptor” partakes in pheromone perception. Similar to osas#9 biosynthesis, *tyra-2* transcript levels are increased in starved animals (Fig. S2C). Notably, octopamine, the distinguishing structural feature of osas#9, has been implicated in responses to food scarcity in invertebrates, including insects (13, 63, 63), *C. elegans* (36, 65–70), and molluscs (71, 72). These findings indicate that worms navigate adverse environmental conditions in part via social communication channels that employ signaling molecules and receptors derived from relevant endocrine signaling pathways.

**Figure 7.**
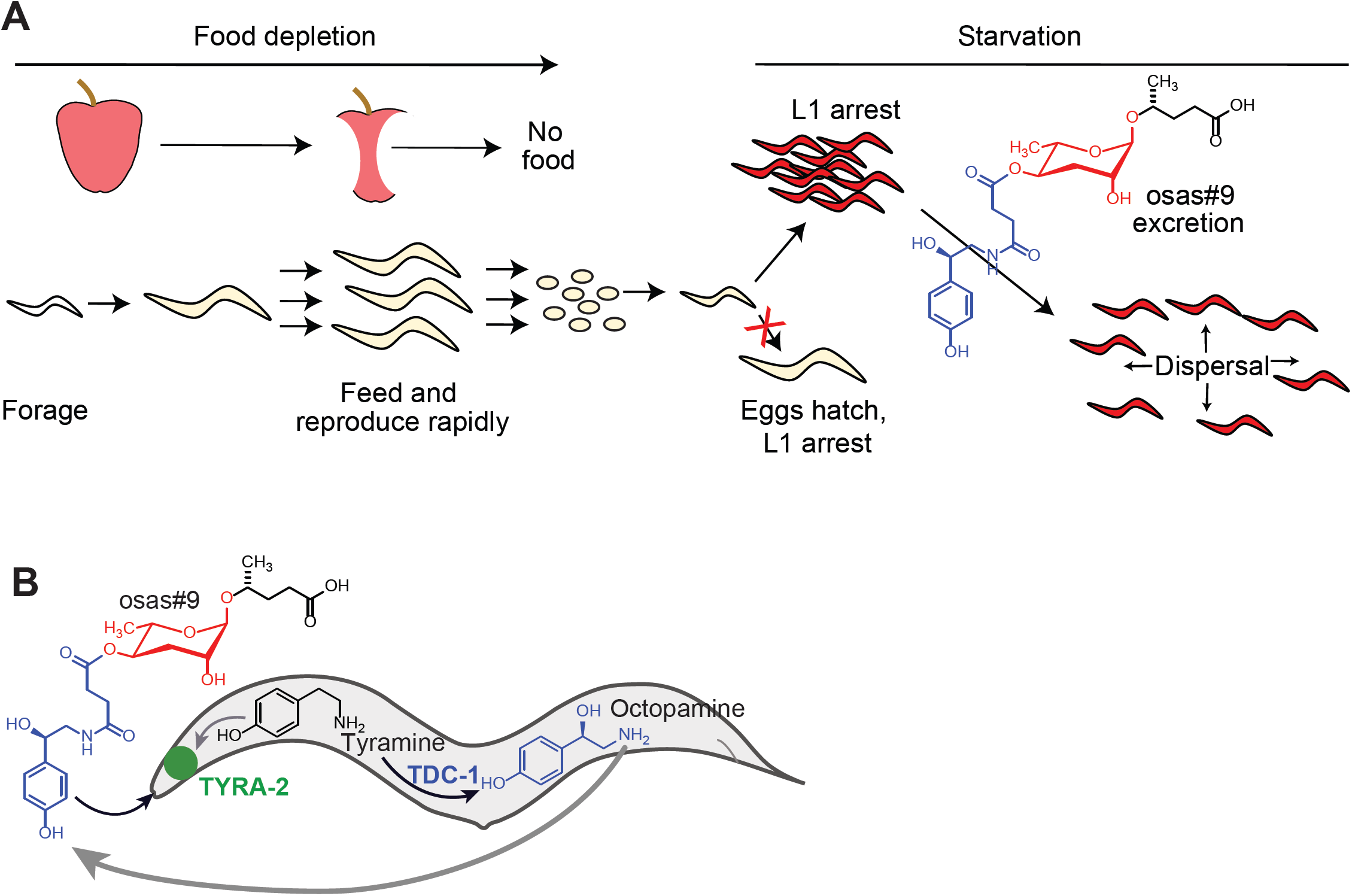
osas#9 serves as a dispersal cue in *C. elegans*. **A)** An animal navigating its environment encounters a food source, and offspring grow and reproduce rapidly, eventually depleting the food. Eggs hatch on depleted food patch and halt development as L1 arrest animals. L1 arrest animals secrete the aversive compound, osas#9 assisting in dispersal away from unfavorable conditions. **B)** Inter-organismal signaling coopts neurotransmitter signaling in *C. elegans*. The G protein-coupled receptor *tyra-2,* which senses tyramine is also required for sensing the biogenic metabolite osas#9 derived from octopamine, to mediate avoidance behavior.

Previous studies have identified several GPCRs involved in ascaroside (ascr) perception: *srbc-64, srbc-66* (ascr#1,2,3) (33); *srg-36, srg-37* (ascr#5) (31); *srx-43, srx-44* (icas#9) (29, 30); *daf-37* (ascr#2), *daf-38* (ascr#2,3,5) (32). These studies demonstrate that GPCRs involved in ascaroside perception may act as heterodimers (32). TYRA-2 has previously been shown to contain the conserved Asp^3.32^ required for amine binding, allowing the receptor to bind tyramine with high affinity, and octopamine to a lesser extent (38). In contrast, osas#9 lacks the basic amine, and instead has an amide as well as an acidic sidechain. These chemical considerations suggest that TYRA-2 may facilitate osas#9 perception by interacting with another GPCR that directly binds to osas#9. However, by ectopically expressing *tyra-2* in ADL and AWA neurons, we were able to elicit responses characteristic to each neuron (Fig. 5). These data show that the response to osas#9 depends on the neuron *tyra-2* is expressed in, providing additional support for direct involvement of TYRA-2 in chemosensation of osas#9. Alternatively, a different receptor that directly interacts with TYRA-2 and is expressed in the ASH, ADL, and AWA neurons could bind osas#9.

Our data suggests that ASE and ASI sensory neurons may regulate ASH sensitivity during osas#9 avoidance serving as modulators at the sensory level, similar to previously observed cross inhibition of ASI and ASH neuronal activity in avoidance to copper, and decision making based on physiological state (73, 74). Alternatively, these neurons could be interacting with ASH neuronal targets in the osas#9 response, strengthening or dampening the relayed signal, possibly through peptidergic or aminergic signaling to establish the functional circuit. Recent studies have shown that *tyra-2* is necessary for binding tyramine in a RIM-ASH feedback loop in multisensory decision making (75). Animals lacking TYRA-2, or the tyramine biosynthetic enzyme TDC-1, crossed a 3M fructose barrier towards an attractant, diacetyl, faster than wildtype *C. elegans.* This demonstrated the endogenous role of tyramine binding to TYRA-2 increasing avoidance in multisensory threat tolerance (75); however, our results show that tyramine signaling is not involved in the response to osas#9. It will be interesting to elucidate the role other neurons or tissues and neuromodulatory signaling have in shaping the osas#9 response. Such modulation of the osas#9 response circuitry remains to be investigated.

Our findings demonstrate that TYRA-2, a member of a well conserved family of neurotransmitter receptors, functions in chemosensation of osas#9, a neurotransmitter-derived inter-organismal signal. Typically, neurotransmitter signaling is intra-organismal, facilitating cell-to-cell communication. This involves the highly regulated biosynthesis of specific chemical compounds, e.g. biogenic amines, their translocation (either by way of diffusion or through active transport), and, finally, perception by dedicated chemoreceptors (76). This mode of communication is strikingly similar to pheromone communication between organisms, as it involves highly specific production and reception of ligands for communication. As evolution is opportunistic, it stands to reason that some machinery from intra-cellular signaling would be utilized for inter-organismal signaling. Indeed, co-option has been hinted at before, in both the trace amine associated receptor (TAAR) and formyl peptide receptor-like (FPRL) receptor classes, both of which are involved in inter-organismal signaling (77–80). Of the TAARs, only TAAR1 and TAAR2 have been found to have endogenous roles: TAAR1 in mammalian CNS, and both TAAR1 and TAAR2 in leukocyte migration (78, 81). Additionally, TAAR2 mRNA has been detected in mouse olfactory epithelium, suggesting it may be involved in both intra-and inter-organismal signaling (77). However, no odor molecules have been linked to TAAR2 in the olfactory epithelium.

How key innovations in metazoan complexity could have evolved from pre-existing machineries is of great interest (82). Our findings demonstrate that the tyramine receptor TYRA-2 functions in chemosensation of osas#9, a neurotransmitter-derived inter-organismal signal, thus revealing involvement of both neurotransmitter biosynthesis and neurotransmitter reception in intra-and inter-organismal signaling. Therefore, evolution of an inter-organismal communication channel co-opted both a small molecule, octopamine, and the related receptor TYRA-2, for mediating starvation-dependent dispersal in *C. elegans* (Fig. 7), suggesting that such co-option may represent one mechanism for the emergence of new inter-organismal communication pathways.

## Methods

### Avoidance drop test

In this assay, the tail end of a forward moving animal is subjected to a small drop (∼5 nl) of solution, delivered through a hand-pulled 10 μl glass capillary tube. The solution, upon contact, is drawn up to the amphid sensory neurons via capillary action. In response, the animal either continues its forward motion (scored as “no avoidance response”), or displays an avoidance response within four seconds (83). The avoidance response is characterized by a reversal consisting of at least one half of a complete “head swing” followed by a change in direction of at least 90 degrees from the original vector. For quantitative analysis, an avoidance response is marked as a “1” and no response as a “0”. The avoidance index is calculated by dividing the number of avoidance responses by the total number of trials. Each trial is done concurrently with osas#9, diluted in DIH_2_O, and a solvent control. Osas#9 was synthesized by methods in Artyukhin et al. 2013 (22).

Integrated mutant strains and controls are prepared using common M9 buffer to wash and transfer a plate of animals to a microcentrifuge tube where the organisms are allowed to settle. The supernatant is removed and the animals are resuspended and allowed to settle again. The supernatant is again removed and the animals then transferred to an unseeded plate. After 1 hour, young adult animals are subjected to the solvent control and the chemical of interest at random with no animal receiving more than one drop of the same solution. Refed animals were transferred to a seeded plate with M9 buffer, and after the allotted time, transferred to an unseeded plate and tested after 10 minutes.

Ablated and extrachromosomal transgenic animals and controls are gently passed onto an unseeded plate and allowed to crawl around. They are then gently passed to another unseeded plate to minimalize bacterial transfer. Ablated animals are tested three times with the solvent control and solution of interest with 2 minute intervals between drops (83).

### Strains and Plasmids

*tyra-2* rescue and misexpression plasmids were generated using MultiSite Gateway Pro Technology and injected into strain FX01846 *tyra-2*(*tm1846*) with co-injection marker p*elt-2*;mCherry by Knudra Transgenics. The promoter attB inserts were generated using PCR and genomic DNA or a plasmid. The *tyra-2* insert was isolated from genomic DNA using attB5*ggcttatccgttgtggagaa* and attB2*ttggcccttccttttctctt.* PDONR221 p1-p5r and PDONR221 P5-P2 donor vectors were used with attB inserts. The resultant entry clones were used with the destination vector pLR305 and pLR306.

### AWA∷tyra-2 misexpression

For AWA expression, a 1.2 kb *odr-10* promoter was isolated from genomic DNA using primers attB1*ctcgctaaccactcggtcat* and attB5r*gtcaactagggtaatccacaattc.* Entry clones were used with destination vector pLR305 resulting in p*odr-10∷tyra-2∷ RFP* and co-injected with p*elt-2*∷mCherry into FX01846.

### ASH∷tyra-2 rescue

For ASH expression, a 3 kb *nhr-79* promoter was isolated from genomic DNA using primers attB1*gtgcaatgcatggaaaattg* and attB5r*atacacttcccacgcaccat.* Entry clones were used with destination vector pLR306 resulting in p*nhr-79∷tyra-2∷RFP* and co-injected with p*elt-2*∷mCherry into FX01846.

### ASH∷*gpa-6* rescue

For ASH expression, a 3 kb *nhr-79* promoter was isolated from genomic DNA using primers attB1*gtgcaatgcatggaaaattg* and attB5r*atacacttcccacgcaccat. gpa-6* was isolated from genomic DNA using primers attB5 cgtctctttcgtttcaggtgtat and attB2 tattttcaaagcgaaacaaaaa. Entry clones were used with destination vector pLR304 resulting in p*nhr-79∷gpa-6∷RFP* and co-injected with p*unc-122*∷RFP into NL1146.

### Translational fusions

*tyra-2∷*GFP fusions were created by PCR fusion using the following primers to isolate 2kb p*tyra-2* with its entire genomic locus from genomic DNA: A) *atgttttcacaagtttcaccaca,* A nested) *ttcacaagtttcaccacattacaa,* and B with overhang) *AGTCGACCTGCAGGCATGCAAGCT gacacgagaagttgagctgggtttc.* GFP primers as described in WormBook (84). The construct was then co-injected with p*elt-2*∷mCherry into both N2 and FX01846.

*gpa-6∷*RFP was generated by adding the restriction sites, AgeI and KpnI, to isolate 4kb p*gpa-6* and the entire *gpa-6* locus from genomic DNA using primers: acatctggtacccctcaatttcccacgatct and acatctaccggtctcatgtaatccagcagacc. RFP∷*unc-54*, ori, and AMPr was isolated from p*unc-122*∷RFP plasmid by PCR addition of the restriction sites AgeI and KpnI with primers: acatctaccggt ATGGTGCGCTCCTCCAAG and ttaataggtaccTGGTCATAGCTGTTTCCTGTG. After digestion and ligation, the clone was injected into N2 with co-injection marker p*unc-122*∷GFP.

(See Supplementary Table 1-3 for details on strains, plasmids, and primers used in this study.)

### RNA interference

RNAi knockdown experiments were performed by following the RNAi feeding protocol found at Source Bioscience (https://www.sourcebioscience.com/products/life-sciences-research/clones/rnai-resources/c-elegans-rnai-collection-ahringer/). The RNAi clones (F01E11.5, F14D12.6, and empty pL4440 vector in HT115) originated from the Vidal Library (85), were generously provided by the Ambros Lab at UMASS Medical School. We observed that RNAi worked best when animals were cultured at 15°C. We used the *nre-1(hd20);lin-15B(hd126)* (VH624) strain for the RNAi studies as it has been previously shown to be sensitive to neuronal RNAi (42, 43).

### Laser ablations

Laser ablations were carried out using DIC optics and the MicroPoint laser system following the procedures as outlined in Fang-Yen *et al.* 2012 (86, 87). Ablated animals were assayed 72 hours later, at the young adult stage. All ablated animals were tested in parallel with control animals that were treated similarly as ablated animals but were not exposed to the laser microbeam.

### Imaging

Translational fusion animals were prepared for imaging by mounting them to a 4% agar pad with 10 mM levamisole on a microscope slide as outlined in O’Hagen and Barr 2016 (88). Animals were imaged using a Nikon Multispectral Multimode Spinning Disk Confocal Microscope, courtesy of Dr. Kwonmoo Lee at Worcester Polytechnic Institute or a Zeiss LSM700 Confocal Microscope, courtesy of the Department of Neurobiology at University of Massachusettes Medical School, Worcester.

Calcium imaging was perfomed by using a modified olfactory chip as described in Reilly et al 2017 (49, 50). A young adult animal was immobilized in a PDMS olfactory chip with its nose subject to a flowing solution. Animals were imaged at 40x magnification for 30 seconds, and experienced a 10 second pulse of osas#9 in between the solvent control. Each animal was exposed to the stimulus three times. Soma fluorescence from GCaMP3 was measured using ImageJ. Background subtraction was performed for each frame to obtain the value ΔF. Change in fluorescence (ΔF/F_0_) was calculated by dividing the ΔF value of each frame by F_0_. F_0_ was calculated as the average ΔF of 10 frames prior to stimulus exposure (50).

### RT-qPCR

RNA was isolated from individual animals, either freshly removed from food or after four hours of starvation using Proteinase K buffer as previously published (89). cDNA was subsequently synthesized using the Maxima H Minus First Strand cDNA Synthesis Kit. iTaq Universal SYBR Green Supermix was used for amplification with the Applied Biosystem 7500 Real Time system. Primer efficiency was determined to be 97.4% for *tyra-2* primers (GAGGAGGAAGAAGATAGCGAAAG, TGTGATCATCTCGCTTTTCA) and 101.8% for the reference gene *ama-1* (GGAGATTAAACGCATGTCAGTG, ATGTCATGCATCTTCCACGA) using the equation 10^(−1/slope)-1. Technical replicates with large standard deviations and trials with a Ct within 5 cycles of the negative control (no reverse transcriptase used in prep) were removed from analyses.

### Locomotion

Speed: Five animals were gently transferred to a 35mm plate and filmed for 20 minutes. Videos were generated using the Wormtracker system by MBF Bioscience. Videos were then analyzed and average speed was computed using software WormLab4.1 (MBF Bioscience, Williston, VT USA).

### Chemoattraction

Diacetyl chemotaxis assays were carried out as previously published, with slight modifications (53). 10 animals were placed in the center of a 35mm plate, equidistant from two spots, one containing 1 µl of solvent control and the other 1 µl of 10^-2^ diacetyl. Both spots contained sodium azide for anesthetizing animals that entered the region. After 45 minutes, the chemotaxis index was calculated by subtracting the number of animals in the solvent control from the number of animals in the solution of interest and divided by the total number of animals.

### Leaving Assay

The leaving assay consisted of the use of 60 mm culture plates containing standard NGM agar. A transparency template that included a 6mm diameter circle in the center was attached to the underside of the NGM plate. One hour before running the assay, young adult animals were passed on to an unseeded plate and allowed to starve for one hour. 100 µl of *E. coli* OP50 liquid culture was spread onto a separate NGM assay plates. These plates were allowed to dry at 25°C without a lid for one hour. After an hour of incubation, 4 µl of either solvent control or 10 pM osas#9 was pipetted onto the agar within the center circle outlined on the template. 10 animals were gently passed into the center circle and their movement was recorded. At one minute intervals, the distance the animals traveled from the origin was measured using ImageJ.

### Statistical analysis

Statistical tests were run using Graphpad Prism. For all figures, when comparing multiple groups, ANOVAs were performed followed by Sidak’s multiple comparison test. When only two groups were compared, a Student’s t-test was used (Figure 1D, S2C). When comparing different strains/conditions, normalized values of osas#9 avoidance index response relative to the respective solvent control were used. This was done to account for any differences in baseline response to solvent control for the respective genotypes, laser ablations, or physiological conditions. When normalizing fold change of osas#9 response to solvent control response for the avoidance assay within a strain/condition, data was first log transformed so a fold change could still be calculated for control plates that had a “0” value. For avoidance assays, statistical groups were based on the number of plates assayed, not the number of drops/animals. For calcium imaging, averages were calculated by obtaining the max peak value before and during exposure to the chemical of interest for each trial.

## Acknowledgements

We thank the *Caenorhabditis* Genetics Center (CGC), which is funded by the NIH Office of Research Infrastructure Programs (P40 OD010440), R. Komuniecki, S. Suo, D. Chase, V. Ambros, C. Bargmann, E.M. Schwarz, and P. Sternberg for strains; R. Garcia, D. Albrecht, and S. Chalasani for plasmids; Knudra transgenics and W. Joyce for injections; K. Lee for the use of the spinning-disk confocal microscope; UMMS Neurobiology department and M. Gorczyca for assistance and use of confocal microscope; V. Ambros, Dana-Farber Cancer Institute, and BioScience Life Sciences for Vidal library RNAi clones; A. Maurya and Piali Sengupta for technical suggestions; D. Vargas Blanco for RT-qPCR guidance; the Srinivasan lab, Rick Komuniecki, Michael Nitabach and Nitabach lab and S. Chalasani for critical comments on the manuscript; A. Warty for contribution to glycerol assays. This work was supported in by grants from the NIH (R01DC016058 to J.S. and GM113692 and GM088290 to FCS and GM084491 to MJA).

## Author Contributions

CDC performed the molecular biology, ablations, and behavioral assays. CDC and LD performed calcium imaging. CDC and VC performed the RNAi behavioral assays. YZ synthesized osas#9. H Choi helped in confocal microscopy of transgenic strains. DR generated strains from MA lab. CDC and JS wrote the manuscript with input from FCS and MJA.

---

**Figure S1.**
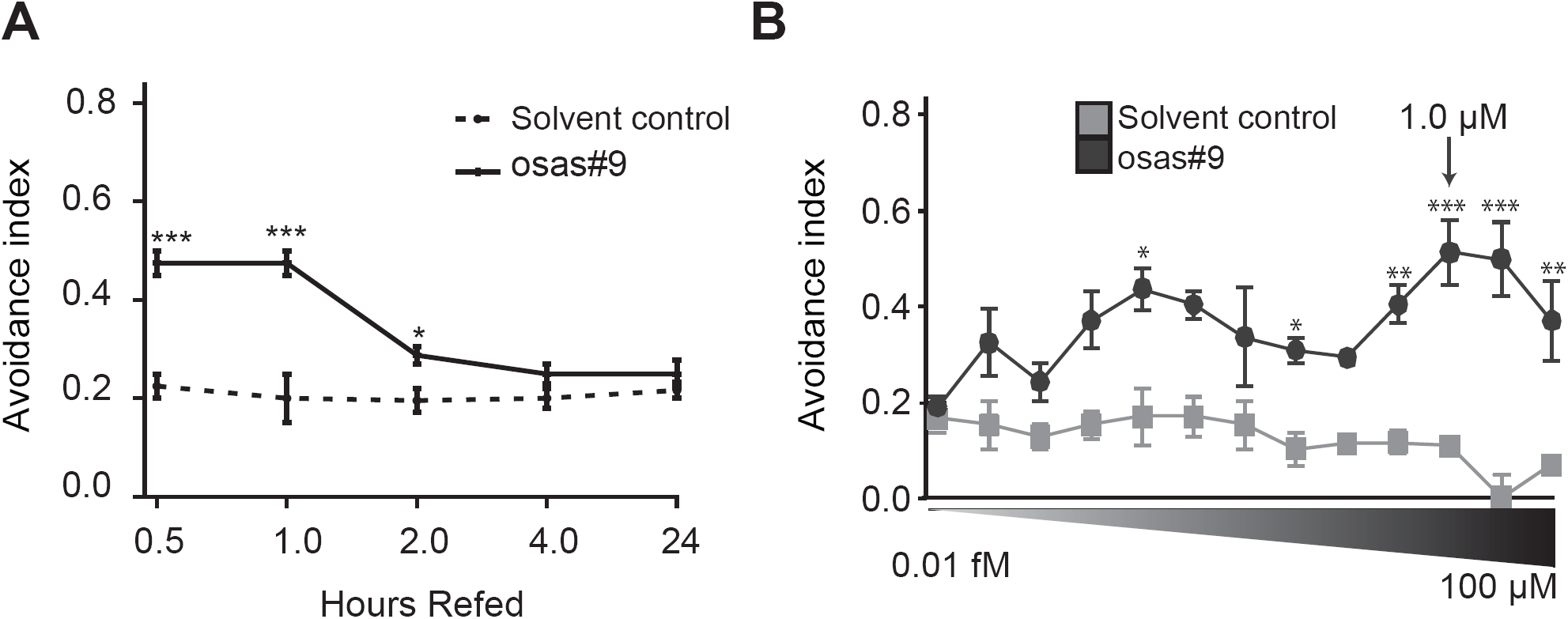
**A)** Attenuation of osas#9 avoidance response by *E. coli* OP50. Animals reintroduced to *E. coli* OP50 for two hours exhibited an attenuated response to osas#9, n≥3 trials. **B)** osas#9 exhibits avoidance response over a broad range of concentrations (fM - µM) in YA wildtype animals, n≥3 trials. Data presented as mean ± S.E.M; *P<0.05, **P<0.01, ***P<0.001, one factor ANOVA with Sidak’s multiple comparison posttest. Asterisks depict comparison between test solution and respective solvent control.

**Figure S2.**
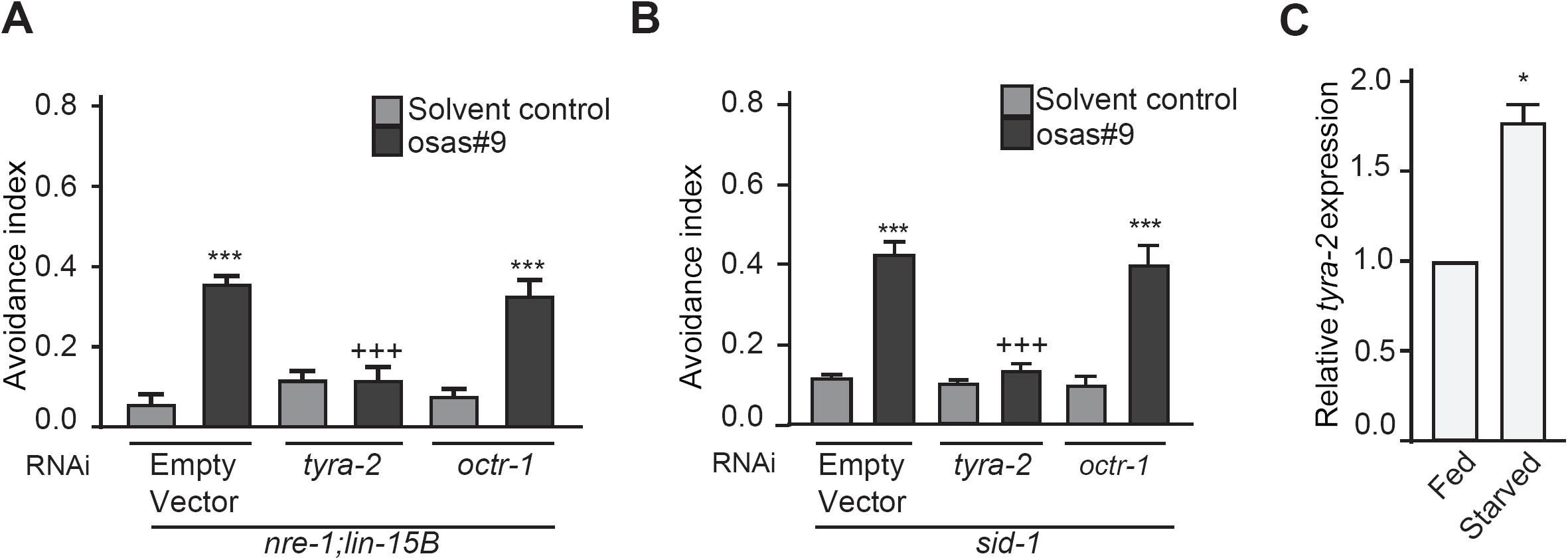
**A-B)** *tyra-2* RNAi knockdown results in loss of avoidance to osas#9. Animals cultured at 15°C and fed *tyra-2* RNAi clones were defective in response to osas#9 in two different RNAi sensitive backgrounds A) *nre-1(hd20) lin-15B(hd126),* n≥10. B) *sid-1(pk3321),* n≥3. **C)** Physiological state dependence of expression of *tyra-2* receptor. RT-qPCR analysis of fed versus starved animals indicates that starved animals upregulate *tyra-2* nearly two-fold. Data shown is the ratio of endogenous *tyra-2* messenger RNA to *ama-1* messenger RNA from three independent RT-qPCR experiments (See materials and methods for more details), n=3. Data presented as mean ± S.E.M;*P<0.05, ***P<0.001, one factor ANOVA with Sidak’s multiple comparison posttest, except for Fig S2C where student’s t-test was used. Asterisks depict comparison between test solution and respective solvent control. ‘+’ signs represent same p value as asterisks but representing difference between osas#9 avoidance of a strain/conditions in comparison to wildtype.

**Figure S3.**
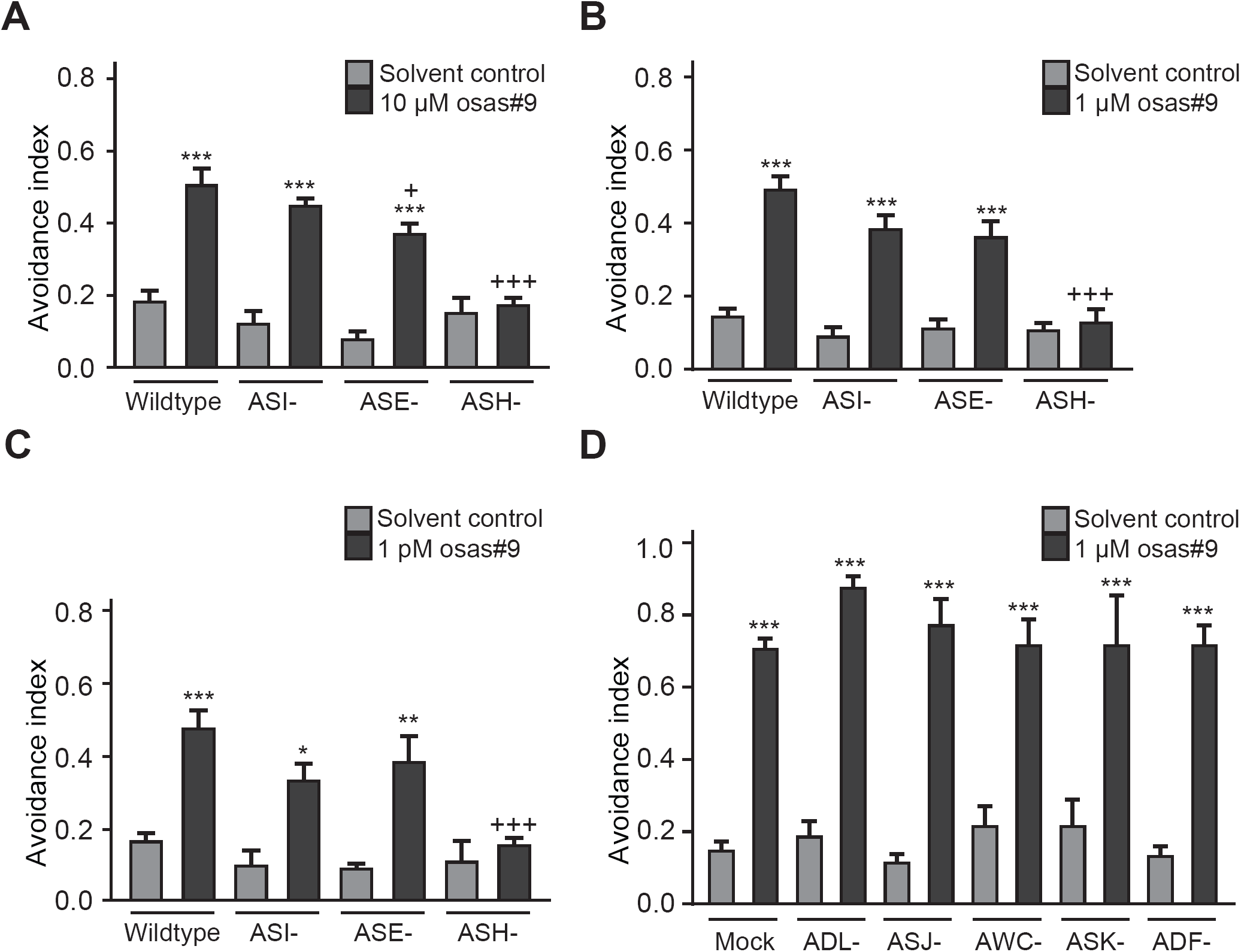
Role of different sensory neurons in osas#9 avoidance behavior. **A-C)** Genetically ablated ASH, ASI and ASE neurons were tested for their response to various concentration of osas#9, n≥3 trials. **D)** Sensory neurons not required for osas#9 avoidance. Note that ADL is not required for osas#9 avoidance. All ablated animals were tested with at least 10 animals with the exception of ADF-, which is 7 animals. Data presented as mean ± S.E.M; *P<0.05, **P<0.01, ***P<0.001, one factor ANOVA with Sidak’s multiple comparison posttest. Asterisks depict comparison between test solution and respective solvent control. ‘+’ signs represent same p value as asterisks but representing difference between osas#9 avoidance of a strain/conditions in comparison to wildtype.

**Figure S4.**
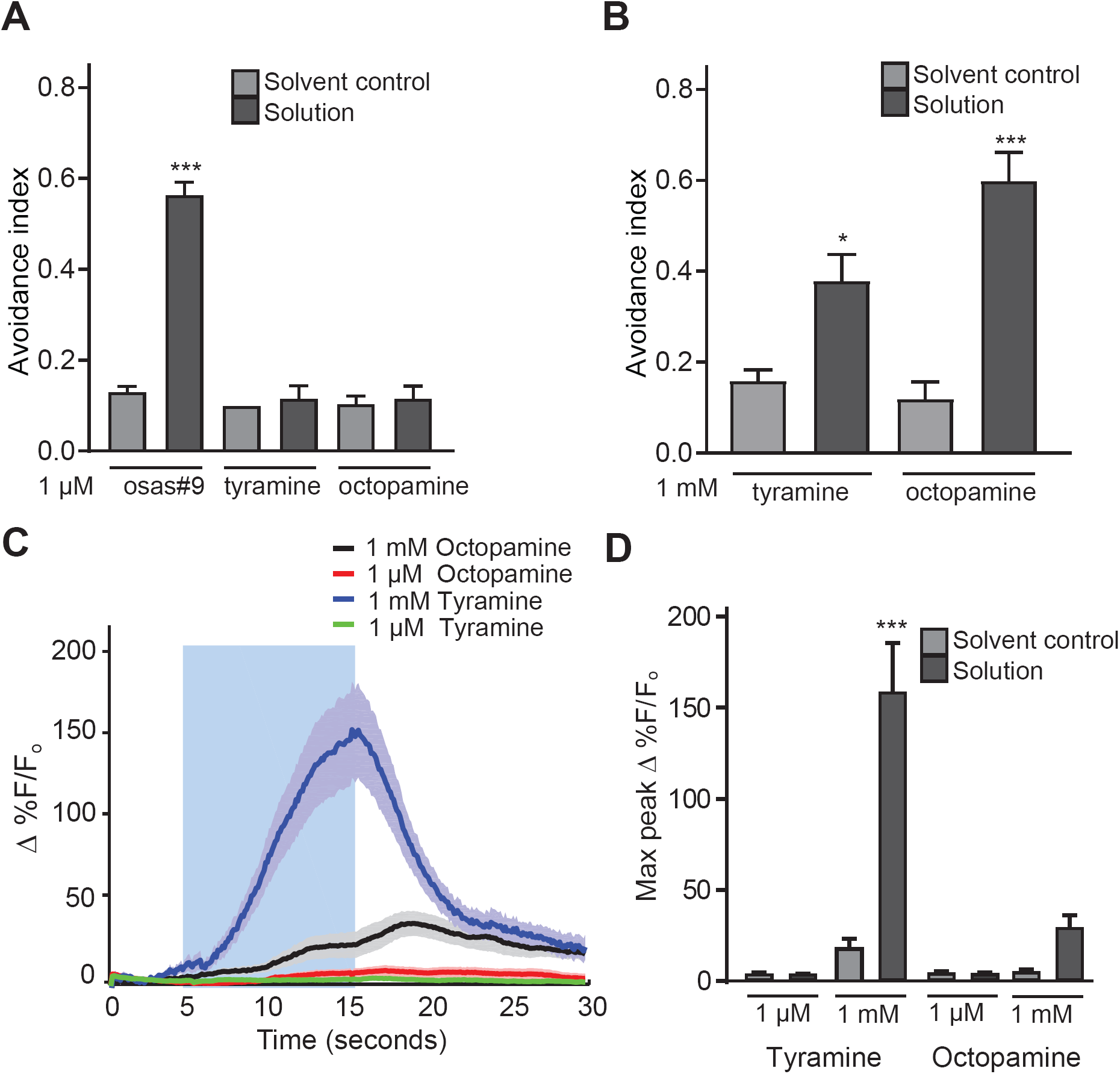
Tyramine and octopamine elicit avoidance at high concentrations. **A)** Animals do not display avoidance to 1 µM tyramine or octopamine, in contrast to osas#9, n≥3 trials. **B)** Tyramine and octopamine result in aversive responses of wildtype animals at higher concentrations, n≥5 trials. **C,D)** Calcium dynamics in ASH sensory neurons upon exposure to tyramine and octopamine. Tyramine exposure resulted in a significant increase in calcium transients in ASH at concentrations of 1 mM, n≥10. Data presented as mean ± S.E.M; *P<0.05, **P<0.01, one factor ANOVA with Sidak’s multiple comparison posttest. Asterisks depict compared solution of interest avoidance response to the solvent control.

**Figure S5.**
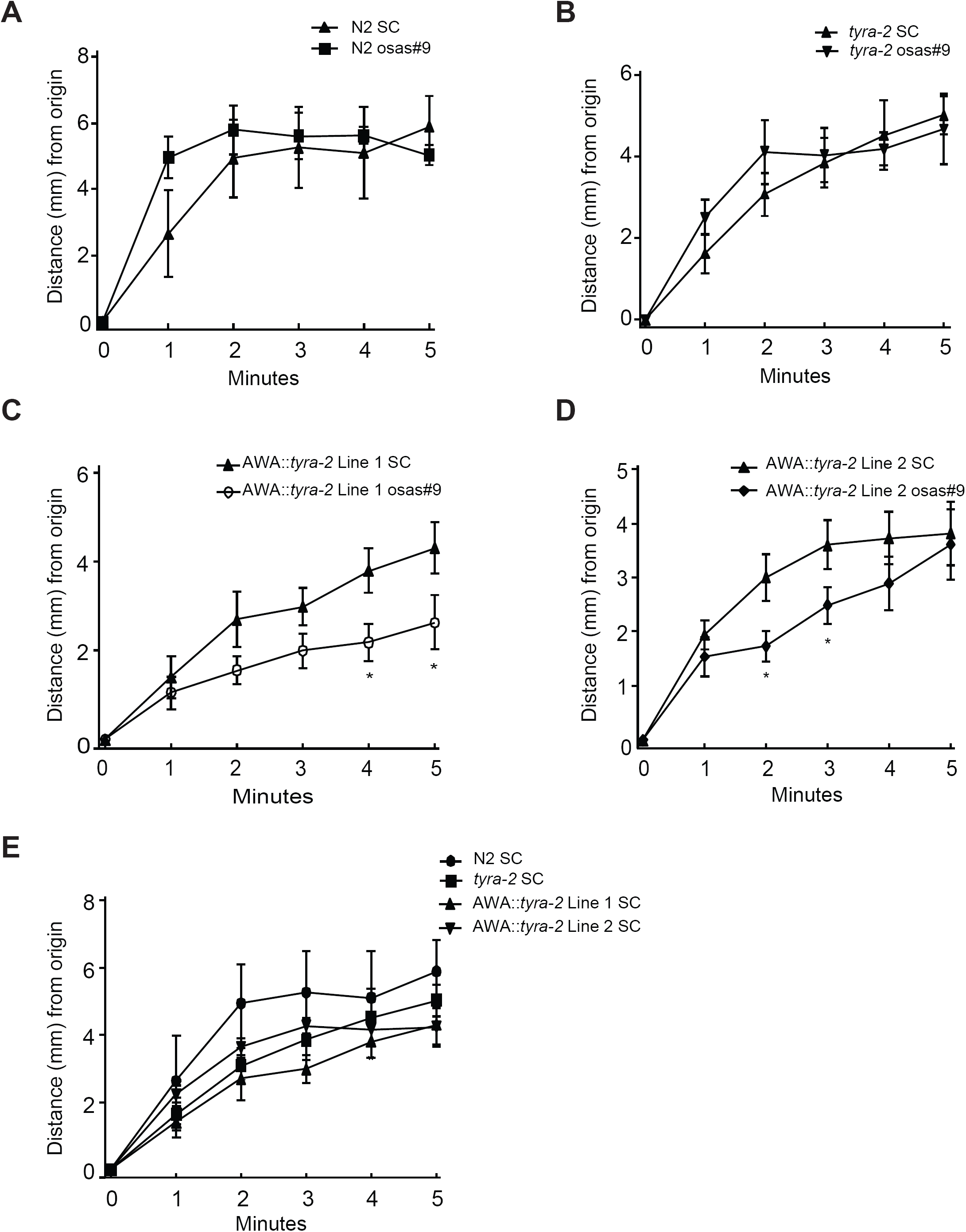
Leaving rates for animals expressing *tyra-2* ectopically in AWA neurons are slower than both wildtype and *tyra-2 lof* animals at 10 pM osas#9. **A)** Wildtype, n≥3 trials. **B)** *tyra-2*, n=6 trials. **C,D)** Two different lines of AWA∷*tyra-2* display slower leaving rates at 10 pM osas#9. n≥6 trials, Line 1 and n≥7 trials, Line 2. **E)** Comparison of solvent control for all strains in leaving assay. None of the animals varied in their response, n≥3 trials. Data presented as mean ± S.E.M; *P<0.05.

**Figure S6.**
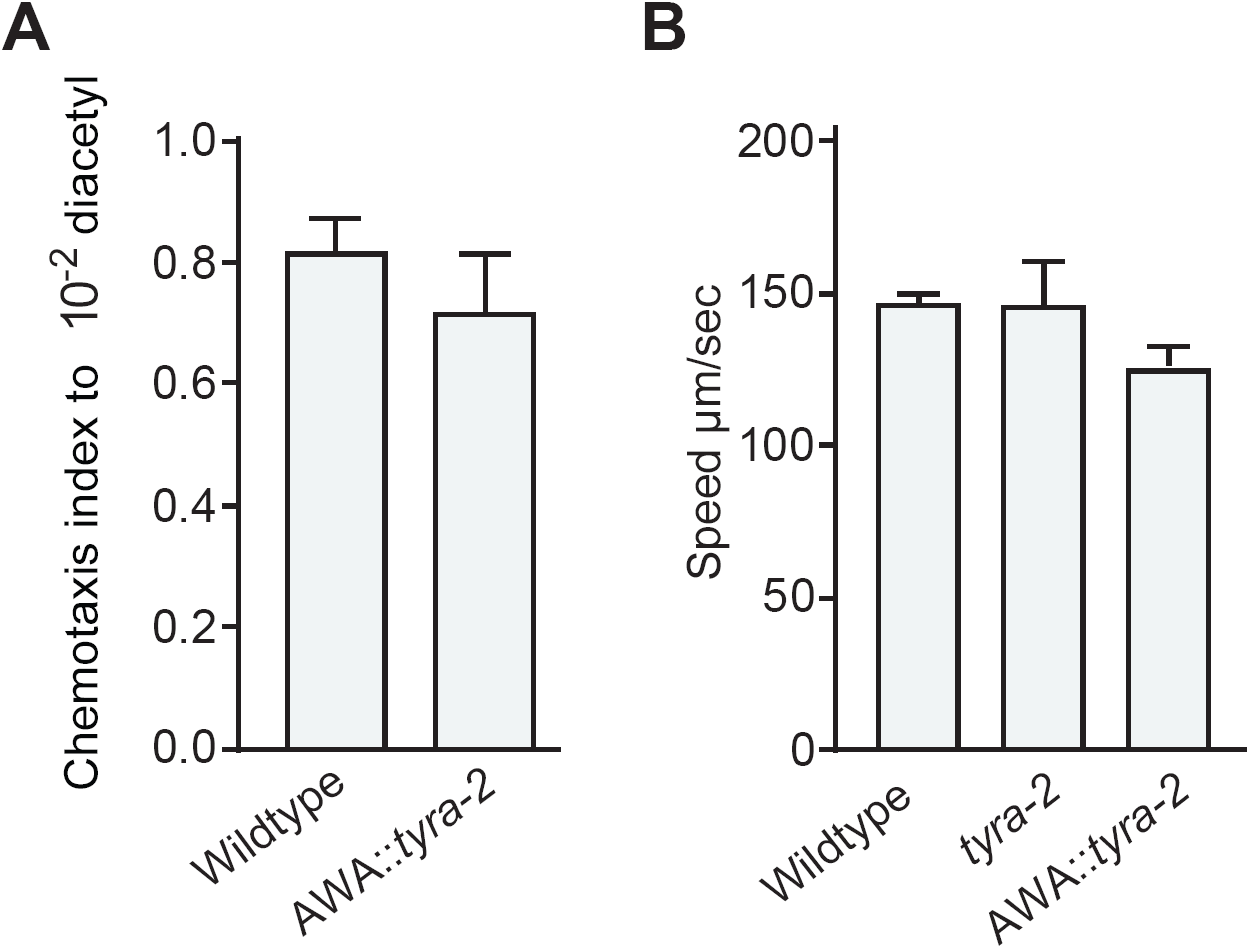
Ectopic expression of *tyra-2* in AWA neurons does not affect AWA-specific behaviors. **A)** Chemotaxis to 10^-2^ diacetyl was unaffected by AWA∷*tyra-2,* n≥7. **B)** Locomotory behaviors were unaltered in AWA∷*tyra-2* animals. Wildtype, *tyra-2 lof*, and AWA∷*tyra-2* speeds are not statistically different, n≥3 trials. Data presented as mean ± S.E.M.

**Video S1.** Video of ASH∷GCaMP3 animal being stimulated with 1 μM osas#9. osas#9 presented to animal when red dot appears on screen. Blue is low level of fluorescence and red is high fluorescence level.

**Table S1.** List of Strains

| Source | Strain | Genotype (allele) | Avoid osas#9? |
| --- | --- | --- | --- |
| Ambros | NL3321 | *sid-1* (pk3321) | yes |
| Alkema | QW42 | *tyra-2* (tm1815) | no |
| Alkema | MT13113 | *tdc-1* (n3419) | yes |
| Alkema | QW569 | *octr-1* (ok371) | yes |
| Alkema | QW284 | *tdc-1* (n3420) | yes |
| Alkema | CX11839 | *tyra-3* (ok325) | yes |
| Alkema | QW1853 | *tyra-2* (tm1846);worEx14[*ptyra-2::tyra-2::GFP @ 1ng/μL*] | yes |
| Bargmann | CX10979 | N2;KyEx2865 [*psra-6::GCAMP3 @ 100 ng/μL*]) | n.d. |
| CGC | CB1489 | *him-8*(e1489) | yes |
| CGC | NL332 | *gpa-1* (pk15)V. | yes |
| CGC | NL335 | *gpa-3* (pk35)V. | yes |
| CGC | NL1146 | *gpa-6* (pk480)X. | no |
| CGC | NL787 | *gpa-11* (pk349)II. | yes |
| CGC | NL2330 | *gpa-13* (pk1270)V. | yes |
| CGC | NL788 | *gpa-14* (pk347)I. | yes |
| CGC | NL797 | *gpa-15* (pk477)I. | yes |
| CGC | CX2205 | *odr-3* (n2150)V. | yes |
| CGC | PR672 | *che-1* (p672) I. | yes |
| Chase | LX702 | *dop-2* (vs105) | yes |
| Chase | LX703 | *dop-3* (vs106) | yes |
| Iino | JN1713 | Is[*sra6p::mCaspl*] | no |
| Komuniecki | FX01846 | *tyra-2* (tm1846) | no |
| Komuniecki | OH313 | *ser-2* (pk1357) | yes |
| Komuniecki | DA1774 | *ser-3* (ad1774) | yes |
| Schwarz | VH624 | rhIs13 [*unc-119::GFP + dpy-20(+)*] V; nre-1(hd20) lin-15B(hd126) X. | yes |
| Srinivasan | JSR19 | *tyra-2* (tm1846);worEx12[pLR306_ *pnhr-79 tyra-2*] | yes |
| Srinivasan | JSR23 | N2;worEx13[*ptyra-2::tyra-2::GFP @ 30ng/μL*] | n.d. |
| Srinivasan | JSR45 | *tyra-2* (tm1846);worEx15[pLR305_ *podr-10 tyra-2*] | no |
| Srinivasan | JSR47 | *tyra-2* (tm1846);worEx15[pLR305_ *podr-10 tyra-2*] | no |
| Srinivasan | JSR50 | *tyra-2* (tm1846) ;KyEx2865 [*psra-6::GCAMP3 @ 100 ng/μL*]) | n.d. |
| Srinivasan | JSR72 | *gpa-6* (pk480)X.; WorEx19 (*pnhr-79::gpa-6::RFP @30ng/ul*; *punc-122::RFP*) | yes |
| Srinivasan | JSR86 | *gpa-6* (pk480)X.; WorEx19 (*pnhr-79::gpa-6::RFP @30ng/ul*; *punc-122::RFP*) | yes |
| Srinivasan | JSR88 | N2; WorEx20 [*pgpa-6::gpa-6::RFP::unc-54 @ 5ng/ul*] | n.d. |
| Sternberg | PY7505 | *oyIs84* [*gpa-4p::TU#813 + gcy-27p::TU#814 + gcy-27p::GFP + unc-122p::DsRed*] | yes |
| Suo | VN280 | *ser-6* (2146) | yes |

**Table S2.**
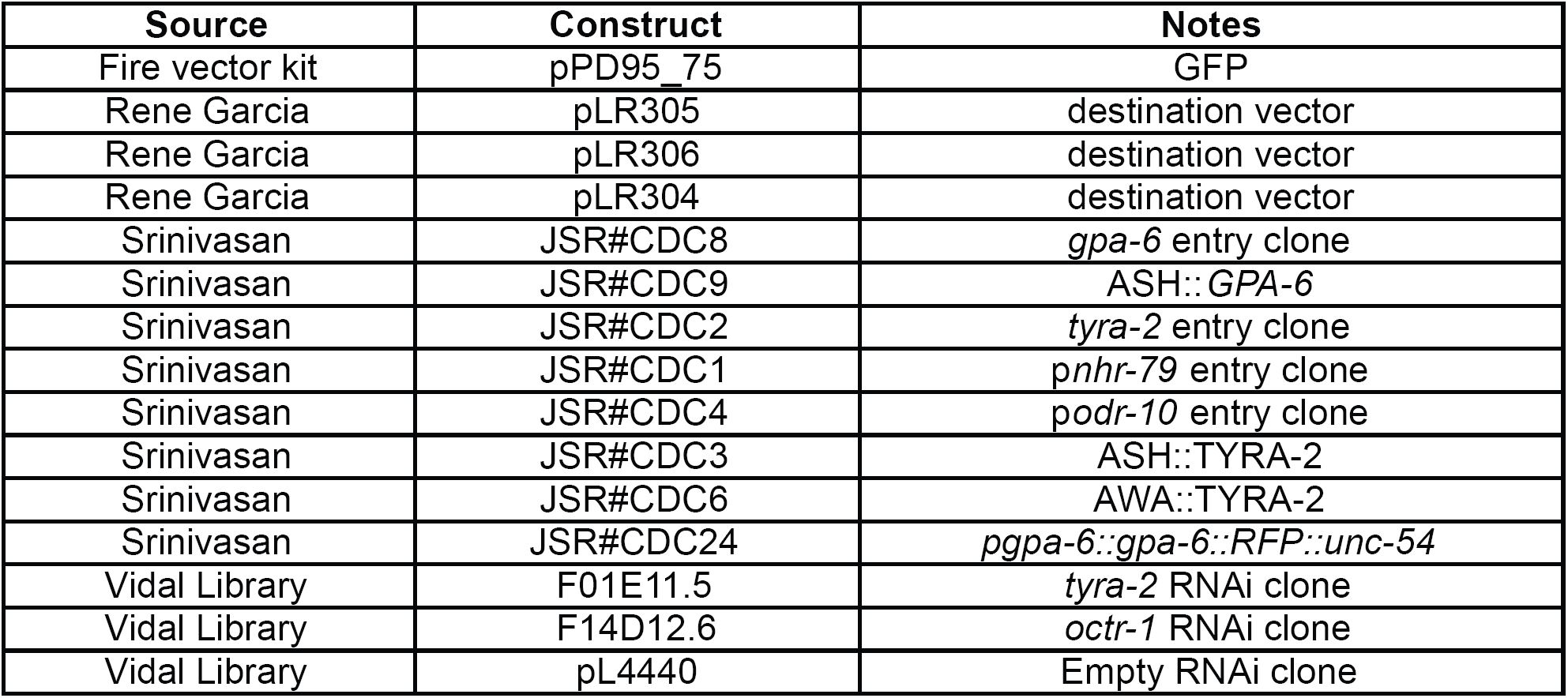
List of Plasmids

| Source | Construct | Notes |
| --- | --- | --- |
| Fire vector kit | pPD95_75 | GFP |
| Rene Garcia | pLR305 | destination vector |
| Rene Garcia | pLR306 | destination vector |
| Rene Garcia | pLR304 | destination vector |
| Srinivasan | JSR#CDC8 | *gpa-6* entry clone |
| Srinivasan | JSR#CDC9 | ASH::*GPA-6* |
| Srinivasan | JSR#CDC2 | *tyra-2* entry clone |
| Srinivasan | JSR#CDC1 | *pnhr-79* entry clone |
| Srinivasan | JSR#CDC4 | *podr-10* entry clone |
| Srinivasan | JSR#CDC3 | ASH::TYRA-2 |
| Srinivasan | JSR#CDC6 | AWA::TYRA-2 |
| Srinivasan | JSR#CDC24 | *pgpa-6::gpa-6::RFP::unc-54* |
| Vidal Library | F01E11.5 | *tyra-2* RNAi clone |
| Vidal Library | F14D12.6 | *octr-1* RNAi clone |
| Vidal Library | pL4440 | Empty RNAi clone |

**Table S3.**
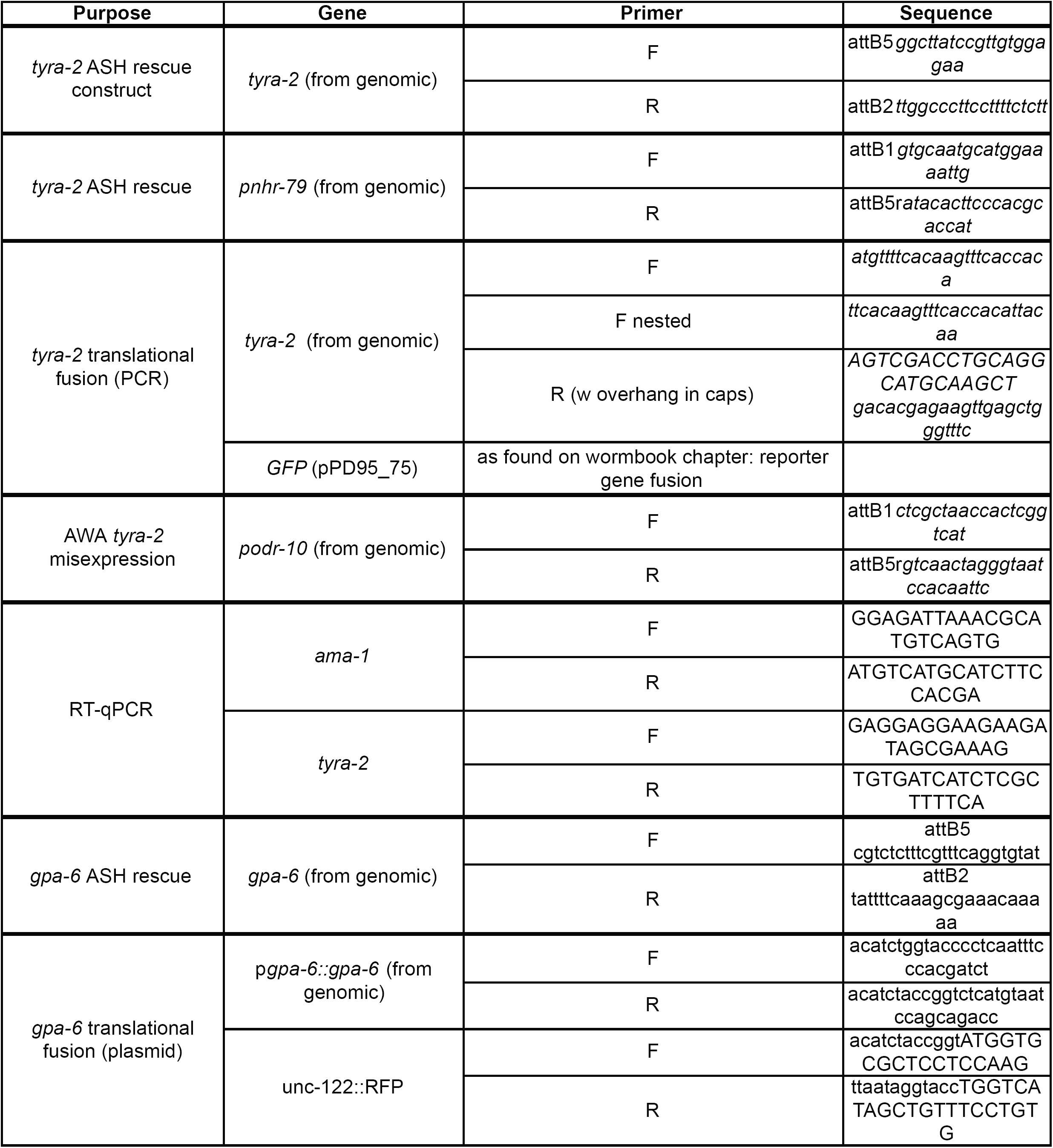
List of Primers

## References

1. Slobodchikoff CN, Paseka A, Verdolin JL. Prairie dog alarm calls encode labels about predator colors. Animal cognition. 2009;12(3):435–9.

2. Barske J, Schlinger BA, Wikelski M, Fusani L. Female choice for male motor skills. Proceedings Biological sciences / The Royal Society. 2011;278(1724):3523–8.

3. Riley JR, Greggers U, Smith AD, Reynolds DR, Menzel R. The flight paths of honeybees recruited by the waggle dance. Nature. 2005;435(7039):205–7.

4. Waters CM, Bassler BL. Quorum sensing: cell-to-cell communication in bacteria. Annual review of cell and developmental biology. 2005;21:319-46.

5. Guerrieri E, Poppy GM, Powell W, Rao R, Pennacchio F. Plant-to-plant communication mediating in-flight orientation of Aphidius ervi. Journal of chemical ecology. 2002;28(9):1703–15.

6. Kunert G, Otto S, Röse USR, Gershenzon J, Weisser WW. Alarm pheromone mediates production of winged dispersal morphs in aphids. Ecology Letters. 2005;8(6):596–603.

7. Akiyama K, Matsuzaki K, Hayashi H. Plant sesquiterpenes induce hyphal branching in arbuscular mycorrhizal fungi. Nature. 2005;435(7043):824–7.

8. Roschina VV. Evolutionary considerations of neurotransmitters in microbial, plant, and animal cells. microbial endocrnology: springer new york; 2010. p. 17-52.

9. Krishnan A, Schioth HB. The role of G protein-coupled receptors in the early evolution of neurotransmission and the nervous system. J Exp Biol. 2015;218(Pt 4):562–71.

10. Berger M, Gray JA, Roth BL. The expanded biology of serotonin. Annu Rev Med. 2009;60:355-66.

11. Chao MY, Komatsu H, Fukuto HS, Dionne HM, Hart AC. Feeding status and serotonin rapidly and reversibly modulate a *Caenorhabditis elegans* chemosensory circuit. Proc Natl Acad Sci U S A. 2004;101(43):15512–7.

12. Tecott LH. Serotonin and the orchestration of energy balance. Cell metabolism. 2007;6(5):352–61.

13. Roeder T. Tyramine and octopamine: ruling behavior and metabolism. Annual review of entomology. 2005;50:447-77.

14. Roeder T. Octopamine in invertebrates. Prog Neurobiol. 1999;59(5):533–61.

15. Gainetdinov RR, Hoener MC, Berry MD. Trace Amines and Their Receptors. Pharmacol Rev. 2018;70(3):549–620.

16. Bargmann CI. Neurobiology of the *Caenorhabditis elegans* genome. Science. 1998;282(5396):2028–33.

17. White JG, Southgate E, Thomson JN, Brenner S. The structure of the nervous system of the nematode *Caenorhabditis elegans*. Philos Trans R Soc Lond B Biol Sci. 1986;314(1165):1–340.

18. Chute CD, Srinivasan J. Chemical mating cues in *C. elegans*. Seminars in cell & developmental biology. 2014;33:18–24.

19. Srinivasan J, Kaplan F, Ajredini R, Zachariah C, Alborn HT, Teal PEA, et al. A blend of small molecules regulates both mating and development in *Caenorhabditis elegans*. Nature. 2008;454(7208):1115–8.

20. Schroeder FC. Modular assembly of primary metabolic building blocks: a chemical language in *C. elegans*. Chem Biol. 2015;22(1):7–16.

21. von Reuss SH, Schroeder FC. Combinatorial chemistry in nematodes: modular assembly of primary metabolism-derived building blocks. Natural product reports. 2015;32(7):994–1006.

22. Artyukhin AB, Yim JJ, Srinivasan J, Izrayelit Y, Bose N, von Reuss SH, et al. Succinylated octopamine ascarosides and a new pathway of biogenic amine metabolism in *Caenorhabditis elegans*. The Journal of biological chemistry. 2013;288(26):18778–83.

23. Kaplan F, Srinivasan J, Mahanti P, Ajredini R,Durak O, Nimalendran R, et al. Ascaroside expression in *Caenorhabditis elegans* is strongly dependent on diet and developmental stage. PLoS One. 2011;6(3):e17804.

24. Butcher RA, Ragains JR, Kim E, Clardy J. A potent dauer pheromone component in *Caenorhabditis elegans* that acts synergistically with other components. Proc Natl Acad Sci U S A. 2008;105(38):14288–92.

25. Jeong PY, Jung M, Yim YH, Kim H, Park M, Hong E, et al. Chemical structure and biological activity of the *Caenorhabditis elegans* dauer-inducing pheromone. Nature. 2005;433(7025):541–5.

26. Srinivasan J, Kaplan F, Ajredini R, Zachariah C, Alborn HT, Teal PE, et al. A blend of small molecules regulates both mating and development in *Caenorhabditis elegans*. Nature. 2008;454(7208):1115–8.

27. Butcher RA, Fujita M, Schroeder FC, Clardy J. Small-molecule pheromones that control dauer development in *Caenorhabditis elegans*. nature chemical biology. 2007;3(7):420–2.

28. Pungaliya C, Srinivasan J, Fox BW, Malik RU, Ludewig AH, Sternberg PW, et al. A shortcut to identifying small molecule signals that regulate behavior and development in *Caenorhabditis elegans*. Proc Natl Acad Sci U S A. 2009;106(19):7708–13.

29. Greene JS, Brown M, Dobosiewicz M, Ishida IG, Macosko EZ, Zhang X, et al. Balancing selection shapes density-dependent foraging behaviour. Nature. 2016;539(7628):254–8.

30. Greene JS, Dobosiewicz M, Butcher RA, McGrath PT, Bargmann CI. Regulatory changes in two chemoreceptor genes contribute to a *Caenorhabditis elegans* QTL for foraging behavior. eLife. 2016;5.

31. McGrath PT, Xu Y, Ailion M, Garrison JL, Butcher RA, Bargmann CI. Parallel evolution of domesticated Caenorhabditis species targets pheromone receptor genes. Nature. 2011;477(7364):321–5.

32. Park D, O’Doherty I, Somvanshi RK, Bethke A, Schroeder FC, Kumar U, et al. Interaction of structure-specific and promiscuous G-protein-coupled receptors mediates small-molecule signaling in *Caenorhabditis elegans*. Proc Natl Acad Sci U S A. 2012;109(25):9917–22.

33. Kim K, Sato K, Shibuya M, Zeiger DM, Butcher RA, Ragains JR, et al. Two chemoreceptors mediate developmental effects of dauer pheromone in *C. elegans*. Science. 2009;326(5955):994–8.

34. Narayan A, Venkatachalam V, Durak O, Reilly DK, Bose N, Schroeder FC, et al. Contrasting responses within a single neuron class enable sex-specific attraction in *Caenorhabditis elegans*. Proc Natl Acad Sci U S A. 2016;113(10):E1392–401.

35. Srinivasan J, von Reuss SH, Bose N, Zaslaver A, Mahanti P, Ho MC, et al. A modular library of small molecule signals regulates social behaviors in *Caenorhabditis elegans*. PLoS biology. 2012;10(1):e1001237.

36. Suo S, Kimura Y, Van Tol HH. Starvation induces cAMP response element-binding protein-dependent gene expression through octopamine-Gq signaling in *Caenorhabditis elegans*. The Journal of neuroscience : the official journal of the Society for Neuroscience. 2006;26(40):10082–90.

37. Mills H, Wragg R, Hapiak V, Castelletto M, Zahratka J, Harris G, et al. Monoamines and neuropeptides interact to inhibit aversive behaviour in *Caenorhabditis elegans*. The EMBO journal. 2012;31(3):667–78.

38. Rex E, Hapiak V, Hobson R, Smith K, Xiao H, Komuniecki R. TYRA-2 (F01E11.5): a *Caenorhabditis elegans* tyramine receptor expressed in the MC and NSM pharyngeal neurons. Journal of neurochemistry. 2005;94(1):181–91.

39. Rex E, Komuniecki RW. Characterization of a tyramine receptor from *Caenorhabditis elegans*. Journal of neurochemistry. 2002;82(6):1352–9.

40. Wragg RT, Hapiak V, Miller SB, Harris GP, Gray J, Komuniecki PR, et al. Tyramine and octopamine independently inhibit serotonin-stimulated aversive behaviors in *Caenorhabditis elegans* through two novel amine receptors. The Journal of neuroscience : the official journal of the Society for Neuroscience. 2007;27(49):13402–12.

41. Calixto A, Chelur D, Topalidou I, Chen X, Chalfie M. Enhanced neuronal RNAi in *C. elegans* using SID-1. Nat Methods. 2010;7(7):554–9.

42. Poole RJ, Bashllari E, Cochella L, Flowers EB, Hobert O. A Genome-Wide RNAi Screen for Factors Involved in Neuronal Specification in *Caenorhabditis elegans*. PLoS Genet. 2011;7(6):e1002109.

43. Schmitz C, Kinge P, Hutter H. Axon guidance genes identified in a large-scale RNAi screen using the RNAi-hypersensitive *Caenorhabditis elegans* strain nre-1(hd20) lin-15b(hd126). Proc Natl Acad Sci U S A. 2007;104(3):834–9.

44. Alkema MJ, Hunter-Ensor M, Ringstad N, Horvitz HR. Tyramine Functions independently of octopamine in the *Caenorhabditis elegans* nervous system. Neuron. 2005;46(2):247–60.

45. Uchida O, Nakano H, Koga M, Ohshima Y. The *C. elegans* che-1 gene encodes a zinc finger transcription factor required for specification of the ASE chemosensory neurons. Development. 2003;130(7):1215–24.

46. Beverly M, Anbil S, Sengupta P. Degeneracy and neuromodulation among thermosensory neurons contribute to robust thermosensory behaviors in *Caenorhabditis elegans*. The Journal of neuroscience : the official journal of the Society for Neuroscience. 2011;31(32):11718–27.

47. Taniguchi G, Uozumi T, Kiriyama K, Kamizaki T, Hirotsu T. Screening of odor-receptor pairs in *Caenorhabditis elegans* reveals different receptors for high and low odor concentrations. Sci Signal. 2014;7(323):ra39.

48. Yoshida K, Hirotsu T, Tagawa T, Oda S, Wakabayashi T, Iino Y, et al. Odour concentration-dependent olfactory preference change in *C. elegans*. Nat Commun. 2012;3:739.

49. Chronis N, Zimmer M, Bargmann CI. Microfluidics for in vivo imaging of neuronal and behavioral activity in *Caenorhabditis elegans*. Nat Methods. 2007;4(9):727–31.

50. Douglas K. Reilly DEL, Dirk R. Albrecht, Jagan Srinivasan. Using an Adapted Microfluidic Olfactory Chip for the Imaging of Neuronal Activity in Response to Pheromones in Male *C. elegans* Head Neurons. JOVE. In Press.

51. Miyabayashi T, Palfreyman MT, Sluder AE, Slack F, Sengupta P. Expression and function of members of a divergent nuclear receptor family in *Caenorhabditis elegans*. Dev Biol. 1999;215(2):314–31.

52. Troemel ER, Kimmel BE, Bargmann CI. Reprogramming chemotaxis responses: sensory neurons define olfactory preferences in *C. elegans*. Cell. 1997;91(2):161–9.

53. Bargmann CI, Hartwieg E, Horvitz HR. Odorant-selective genes and neurons mediate olfaction in *C. elegans*. Cell. 1993;74(3):515–27.

54. Sengupta P, Chou JH, Bargmann CI. odr-10 encodes a seven transmembrane domain olfactory receptor required for responses to the odorant diacetyl. Cell. 1996;84(6):899–909.

55. Troemel ER, Chou JH, Dwyer ND, Colbert HA, Bargmann CI. Divergent seven transmembrane receptors are candidate chemosensory receptors in *C. elegans*. Cell. 1995;83(2):207–18.

56. Sambongi Y, Nagae T, Liu Y, Yoshimizu T, Takeda K, Wada Y, et al. Sensing of cadmium and copper ions by externally exposed ADL, ASE, and ASH neurons elicits avoidance response in *Caenorhabditis elegans*. Neuroreport. 1999;10(4):753–7.

57. de Bono M, Tobin DM, Davis MW, Avery L, Bargmann CI. Social feeding in *Caenorhabditis elegans* is induced by neurons that detect aversive stimuli. Nature. 2002;419(6910):899–903.

58. Jang H, Kim K, Neal SJ, Macosko E, Kim D, Butcher RA, et al. Neuromodulatory state and sex specify alternative behaviors through antagonistic synaptic pathways in *C. elegans*. Neuron. 2012;75(4):585–92.

59. Jansen G, Thijssen KL, Werner P, van der Horst M, Hazendonk E, Plasterk RH. The complete family of genes encoding G proteins of *Caenorhabditis elegans*. Nature genetics. 1999;21(4):414–9.

60. Lans H, Rademakers S, Jansen G. A network of stimulatory and inhibitory Galpha-subunits regulates olfaction in *Caenorhabditis elegans*. Genetics. 2004;167(4):1677–87.

61. Bastiani C, Mendel J. Heterotrimeric G proteins in C. elegans. WormBook. 2006:1-25.

62. Felix MA, Braendle C. The natural history of *Caenorhabditis elegans*. Curr Biol. 2010;20(22):R965–9.

63. Li Y, Tiedemann L, von Frieling J, Nolte S, El-Kholy S, Stephano F, et al. The Role of Monoaminergic Neurotransmission for Metabolic Control in the Fruit Fly Drosophila Melanogaster. Front Syst Neurosci. 2017;11:60.

64. Li Y, Hoffmann J, Li Y, Stephano F, Bruchhaus I, Fink C, et al. Octopamine controls starvation resistance, life span and metabolic traits in Drosophila. Scientific reports. 2016;6:35359.

65. Yoshida M, Oami E, Wang M, Ishiura S, Suo S. Nonredundant function of two highly homologous octopamine receptors in food-deprivation-mediated signaling in *Caenorhabditis elegans*. Journal of neuroscience research. 2014;92(5):671–8.

66. Greer ER, Perez CL, Van Gilst MR, Lee BH, Ashrafi K. Neural and molecular dissection of a *C. elegans* sensory circuit that regulates fat and feeding. Cell metabolism. 2008;8(2):118–31.

67. Churgin MA, McCloskey RJ, Peters E, Fang-Yen C. Antagonistic Serotonergic and Octopaminergic Neural Circuits Mediate Food-Dependent Locomotory Behavior in *Caenorhabditis elegans*. The Journal of neuroscience : the official journal of the Society for Neuroscience. 2017;37(33):7811–23.

68. Hoshikawa H, Uno M, Honjoh S, Nishida E. Octopamine enhances oxidative stress resistance through the fasting-responsive transcription factor DAF-16/FOXO in *C. elegans*. Genes Cells. 2017;22(2):210–9.

69. Tao J, Ma YC, Yang ZS, Zou CG, Zhang KQ. Octopamine connects nutrient cues to lipid metabolism upon nutrient deprivation. Sci Adv. 2016;2(5):e1501372.

70. Horvitz HR, Chalfie M, Trent C, Sulston JE, Evans PD. Serotonin and octopamine in the nematode *Caenorhabditis elegans*. Science. 1982;216(4549):1012–4.

71. Aonuma H, Kaneda M, Hatakeyama D, Watanabe T, Lukowiak K, Ito E. Weak involvement of octopamine in aversive taste learning in a snail. Neurobiol Learn Mem. 2017;141:189–98.

72. Vehovszky A, Szabo H, Elliott CJ. Octopamine increases the excitability of neurons in the snail feeding system by modulation of inward sodium current but not outward potassium currents. BMC neuroscience. 2005;6:70.

73. Guo M, Wu TH, Song YX, Ge MH, Su CM, Niu WP, et al. Reciprocal inhibition between sensory ASH and ASI neurons modulates nociception and avoidance in *Caenorhabditis elegans*. Nat Commun. 2015;6:5655.

74. Davis KC, Choi YI, Kim J, You YJ. Satiety behavior is regulated by ASI/ASH reciprocal antagonism. Scientific reports. 2018;8(1):6918.

75. Ghosh DD, Sanders T, Hong S, McCurdy LY, Chase DL, Cohen N, et al. Neural Architecture of Hunger-Dependent Multisensory Decision Making in *C. elegans*. Neuron. 2016;92(5):1049–62.

76. Chase DL, Koelle MR. Biogenic amine neurotransmitters in C. elegans. WormBook. 2007:1–15.

77. Liberles SD, Buck LB. A second class of chemosensory receptors in the olfactory epithelium. Nature. 2006;442(7103):645–50.

78. Babusyte A, Kotthoff M, Fiedler J, Krautwurst D. Biogenic amines activate blood leukocytes via trace amine-associated receptors TAAR1 and TAAR2. Journal of leukocyte biology. 2013;93(3):387–94.

79. Riviere S, Challet L, Fluegge D, Spehr M, Rodriguez I. Formyl peptide receptor-like proteins are a novel family of vomeronasal chemosensors. Nature. 2009;459(7246):574–7.

80. Stempel H, Jung M, Perez-Gomez A, Leinders-Zufall T, Zufall F, Bufe B. Strain-specific Loss of Formyl Peptide Receptor 3 in the Murine Vomeronasal and Immune Systems. The Journal of biological chemistry. 2016;291(18):9762–75.

81. Liberles SD. Trace amine-associated receptors: ligands, neural circuits, and behaviors. Current opinion in neurobiology. 2015;34:1–7.

82. de Mendoza A, Sebe-Pedros A, Ruiz-Trillo I. The evolution of the GPCR signaling system in eukaryotes: modularity, conservation, and the transition to metazoan multicellularity. Genome Biol Evol. 2014;6(3):606–19.

83. Hilliard MA, Bargmann CI, Bazzicalupo P. *C. elegans* responds to chemical repellents by integrating sensory inputs from the head and the tail. Curr Biol. 2002;12(9):730–4.

84. Boulin T, Etchberger JF, Hobert O. Reporter gene fusions. WormBook. 2006:1–23.

85. Rual JF, Ceron J, Koreth J, Hao T, Nicot AS, Hirozane-Kishikawa T, et al. Toward improving *Caenorhabditis elegans* phenome mapping with an ORFeome-based RNAi library. Genome research. 2004;14(10B):2162–8.

86. Fang-Yen C, Gabel CV, Samuel AD, Bargmann CI, Avery L. Laser microsurgery in *Caenorhabditis elegans*. Methods Cell Biol. 2012;107:177–206.

87. Srinivasan J, Durak O, Sternberg PW. Evolution of a polymodal sensory response network. BMC biology. 2008;6:52.

88. O’Hagan R, Barr MM. Kymographic Analysis of Transport in an Individual Neuronal Sensory Cilium in *Caenorhabditis elegans*. Methods in molecular biology. 2016;1454:107–22.

89. Ly K, Reid SJ, Snell RG. Rapid RNA analysis of individual *Caenorhabditis elegans*. MethodsX. 2015;2:59–63.

